# Data-Driven Network Models for Genetic Circuits From Time-Series Data with Incomplete Measurements

**DOI:** 10.1101/2021.03.10.434835

**Authors:** Enoch Yeung, Jongmin Kim, Ye Yuan, Jorge Gonçalves, Richard M. Murray

## Abstract

Synthetic gene networks are frequently conceptualized and visualized as static graphs. This view of biological programming stands in stark contrast to the transient nature of biomolecular interaction, which is frequently enacted by labile molecules that are often unmeasured. Thus, the network topology and dynamics of synthetic gene networks can be difficult to verify *in vivo* or *in vitro*, due to the presence of unmeasured biological states. Here we introduce the dynamical structure function as a new mesoscopic, data-driven class of models to describe gene networks with incomplete measurements. We introduce a network reconstruction algorithm and a code base for reconstructing the dynamical structure function from data, to enable discovery and visualization of graphical relationships in a genetic circuit diagram as *time-dependent functions* rather than static, unknown weights. We prove a theorem, showing that dynamical structure functions can provide a data-driven estimate of the size of crosstalk fluctuations from an idealized model. We illustrate this idea with numerical examples. Finally, we show how data-driven estimation of dynamical structure functions can explain failure modes in two experimentally implemented genetic circuits, a historical genetic circuit and a new *E. coli* based transcriptional event detector.

## I. Introduction

Synthetic gene networks fulfill diverse roles in realizing circuit logic [1] and timing in living organisms [2]. Ranging from single-input inverters [3], [4] to combinatorial input logic gates [5], [6], reduction in DNA synthesis and sequencing costs have made it possible to build increasingly complex genetic circuits with tens to hundreds of components. However, the ability to build novel biological circuitry often outpaces our ability to revise designs or to take verify what has been built behaves as intended. As the fields of synthetic and systems biology continue to build and integrate on successes of circuit and device-level complexity to engineer entire genetic systems or pathways, we are consistently seeing failure modes that arise from a lack of modularity, retroactivity [7], [8], [9], and context effects [10].

Likewise, the expansion of CRISPR-based methods for genome editing and targeted gene knockdown [11], [12] has enabled a broader category of systems biology design problems, centered on redesigning genomes [13] or reprogramming host regulatory networks [14] to target specific environmental niches or to exhibit a particular phenotype. The underlying genetic program implicit in these systems biology objectives is often a vast, complex, and dynamic network of interacting genes, mRNA, and proteins. The expansion in DNA sequencing read depth has made it possible to profile individual genes via the transcriptome [15], which combined with quantitative proteomics [16] or metabolomics [17], enables systems-level analysis of network activity. But prohibitive sampling and library preparation costs make obtaining highly time-resolved omics’ measurements hard. This makes it difficult to infer dynamic network activity at the scale of whole cell models [18] without extensive experimental investment.

Dynamic network models that describe the intricate interactions between every biomolecular state or species are referred to as state-space models. Two key variables that often determine the behavior of these network models are its network topology [19], [20] and parametric realization [21], [22]. The structure of a network is generally determined by how states in the system causally affect each other [22]; edges in the network are determined by causal dependence while nodes are determined by the states of the system [23].

Identifying the active, dynamic network structure of a biological network is critical, since the hypothesized network architecture of a genetic circuit may be very different from the realized network architecture using a specific collection of parts, sequences, and composition approach [24]. While network structure alone does not determine dynamical behavior, though, parametric information is also important in determining what dynamical behaviors a system can achieve [25]. Rather, network structure, or topology, often defines or narrows the possible behaviors a system can achieve. Without any structural constraints, a dynamical system can have arbitrary input-output behavior. Once network structure is imposed, the set of realizable input-output trajectories can be reduced [26], [27]. If the realized network differs significantly from the intended network design, the dynamics of the system may produce faults or glitches when appropriately excited or interrogated [28], [29]. Getting the actual network topology to match the intended network motif is thus a key element to robust synthetic biological design.

In systems and synthetic biology, canonical network motifs are broadly accepted as enabling useful dynamical behavior [27], [30]. For example, an incoherent feedforward loop can be used for fold-change detection or adaptation [31], [32]. A cyclic network of repressors is associated with either oscillations [33], [34], [35] or multi-stability [36] while a dual negative feedback network of two nodes is used as memory module or toggle switch [37]. Still, the active, dynamic network architecture of most realizations of these network motifs in the form of genetic circuits are not formally characterized or catalogued [38]. Systematic, generalizable tools that can discover and model dynamic network topology from data are valuable. [1].

Circuit network discovery is, at its core, a network reconstruction problem. Given a desired network motif and a physical system, we need to use measurements of the system to determine if the actual, active network of the system matches the intended design. There have been many network reconstruction algorithms developed for natural and synthetic biological networks [39], [40], [24], [41], [42], [43], [44], [45]. Historically, the approach to discovering network interactions has involved direct perturbation of biochemical species or components in a network [41], [45], [46]. Individual nodes are perturbed and depending on if nearby nodes positively or negatively correlate, an activating or repressing relationship between two network nodes can be inferred. In [24], this framework was taken a step further, by showing that the behavior of direct and indirect links in a benchmark circuit is network topology dependent. This provided a means for using steady state perturbation data [39], [47], [45] to estimate network models. Further, these steady-state estimation algorithms have been verified using a benchmark synthetic gene cirucit [43]. More recently, the authors in [42] and [40] showed that retroactivity in gene networks can paradoxically confound network predictions that are based wholly on correlation measures. The core issue is that even when measurement data for all biological states is available, causality is difficult to determine from steady-state measurement data affected by back-action or retroactivity in genetic networks [40].

At the single cell level, the reconstruction problem for biological networks introduces challenges of inferring non-linear stochastic models from noisy data [48], [49], [44]. In [48], the authors show that by comparing average abundances, molecule lifetimes, covariances, and magnitude of step, they can map pairwise interaction dynamics, even when the rest of the system is completely unspecified. The key observation is that assembly stoichiometry of new molecules is fixed, so unbalanced production of linked precursor components will exacerbate imbalance further, resulting in empirically observed large fluctuations. Further, Hilfinger, Norman and Paulsson show in [49] that there are statistical invariants for certain kinds of network interactions, which can be used to evaluate and challenge existing hypotheses of stochastic gene interaction. More recently, Wang, Lin, Sontag and Sorger show that effective stoichiometric spaces can be used to determine network structure from the covariances of single-cell multiplex data [44]. These studies show that it is possible to infer meaningful structural information about a genetic network, even only when a portion of the network states are observed and the data is fundamentally noisy.

In this paper we introduce a class of mesoscopic network reconstruction models with adaptable resolution, commensurate with the depth or coverage of the circuit states (or genome) available from fluorimetric, spectometry-based, or sequencing based measurements. Our method is distinct in that we consider the use of high-resolution time-series data, but where only partial measurement of the network’s nodes is feasible. Further, we consider dynamic measurements of bulk culture rather than single cell, where we benefit from the assumptions of high molecular copy number and large reaction volumes [49]. Specifically, we present the dynamical structure function, an abstract model class from linear time-invariant systems theory and show it can be used as a generalized representation of measured interactions between biological or biochemical states. The contributions of this paper are: 1) we show how a dynamical structure function can encode both direct and crosstalk network interactions, by way of theorem and simulated examples, 2) we develop a direct estimation algorithm and code to directly estimate the dynamical structure function, as well as visualization tools to monitor repression and activation in genetic circuits, and 3) we demonstrate this theory on two experimental systems: A) an *in vitro* genelet repressilator from the synthetic biology literature and B) a novel transcriptional event detector that we build specifically to illustrate dynamical structure reconstruction.

## II. Representing Network Interactions in Partially Measured Biological Networks with Dynamical Structure Functions

The tradeoffs between cost of network reconstruction and the “informativity” of the structural representation are especially clear in synthetic and systems biology research. In this area, finding or verifying the network of a biological system is an important problem. However, discovering the entire chemical reaction network is typically an ill-posed problem, since additional reactions may be introduced due to host or environmental context [52], loading effects, or unanticipated retroactivity effects [53], [54], [55], [56], [7]. Even without these effects, the reconstruction problem is equivalent to finding a unique realization for the dynamical system from direct measurements of every state of the system. Unique realization problems are difficult, unless the system of interest has specific structure, e.g., measurement functions of the state that are diffeomorphic [57], [58]. On the other hand, there are many inputs that can be used to perturb the system of interest, e.g. silencing RNA [59], genetic knock-outs [60], and small chemical inducers [61]. Using these inputs, it is straightforward to reconstruct the transfer function of the system [73], [51]. However, the transfer function contains virtually no information about how chemical species within the system are interacting.

An intermediate representation of network structure that addresses this trade-off is the dynamical structure function [62]. The dynamical structure function is a representation derived from linear systems theory, and thus can be used to model transients of a genetic circuit around an operating point or even unstable network dynamics diverging from an equilibrium point. It is a more detailed description of network structure than the transfer function since it models the causal interactions between measured outputs, in addition to the causal dependencies of outputs on input variables. At the same time, it does not require complete state feedback for reconstruction, since it only models the interactions among output states. In biological systems, this is especially applicable since the output variables of a system are also a subset of the state variables. All unmeasured states are subsumed in the edge-weight functions that describe interactions between measured variables. It is thus possible to experimentally target specific biochemical species to measure and verify that the network structure of a biological system is functioning as intended. Most notably, necessary and sufficient conditions for recovery of dynamical network models have been developed and well-studied [62], [22], [63], [64], [65], [66], but so far no open-source algorithms, code bases, or applications of this theory have been developed directly for synthetic biology.

### A. Dynamical Structure Functions

We briefly review the theory of dynamical structure functions, as they pertain to biochemical reaction networks. In practice, the state of the dynamical system 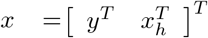 ∈ ℝ^*n*^, where *y* ∈ ℝ^*p*^ are the measured chemical states of the dynamical system, corresponding to components of the biochemical reaction network tagged with fluorescent reporters, and *x*_*h*_ ∈ ℝ^*n*−*p*^ are the unmeasured chemical states. It is also the cases that there are exogenous inputs *u* ∈ ℝ^*m*^ that can be introduced to influence the dynamics of the state *x*. With the exception of oscillators, many biochemical reaction networks converge to a steady state. Moreover, it is generally the case that the parameters of biochemical reaction networks are time-invariant, so long as macroscopic experimental settings of the system such as temperature, growth media, and dissolved oxygen content remain fixed. Therefore, while the model of a biochemical reaction network is of the form

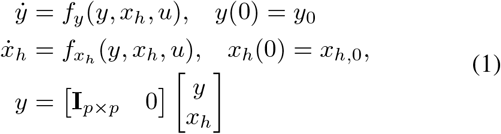

we will suppose that we can linearize the system about either an equilibrium point, a nominal operating point, or even an (unstable or oscillatory) initial condition to extract network dynamics. In biological systems, networks are almost never precisely linear, but we presume to model local fluctuations or perturbations from a target point in the state space. As we will see in the sequel, this will be enough to extract relevant network information. Proceeding with the linearization, we can write the system in the form:

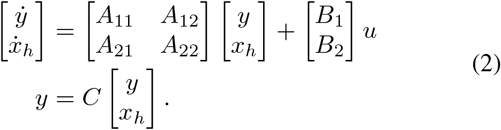

where

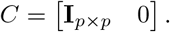

We also assume the system’s initial condition of the linearized system is *x*(0) = 0, and the entries in *A* ∈ ℝ^*n×n*^ and *B* ∈ ℝ^*n×m*^ are calculated as

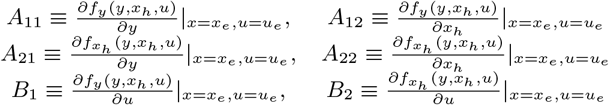

Taking Laplace transforms, solving for *X*_*h*_(*s*) and replacing it in *Y* (*s*) we obtain

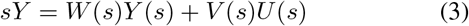

where

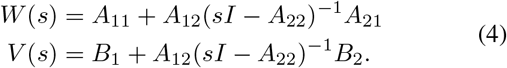

Defining *D*(*s*) = diag (*W* (*s*)) and subtracting *D*(*s*)*Y* (*s*) from both sides of equation (3) and solving for *Y* (*s*) we obtain the following equation

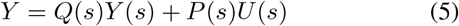

where *Q*(*s*) = (*sI* −*D*)^−1^(*W* −*D*) is a *p* × *p* transfer function matrix and *P* (*s*) = (*sI* − *D*)^−1^*V* is a *p* ×*m* transfer function matrix. Each entry *Q*_*ij*_(*s*) is a transfer function that describes the causal dependency of measured state *Y*_*i*_(*s*) on measured state *Y*_*j*_(*s*). Similarly, the transfer function *P*_*ij*_(*s*) describes the causal dependency of measured state *Y*_*i*_(*s*) on input *U*_*j*_(*s*). The matrix pair (*Q*(*s*), *P* (*s*)) is known as the *dynamical structure function*, where *Q*(*s*) is referred to as the network structure and *P* (*s*) as the control structure.

Note that *Q*(*s*) is defined as *Q*(*s*) = (*sI* −*D*)^−1^(*W* −*D*) rather than *Q*(*s*) = 1*/sW* (*s*). This guarantees that the diagonal entries of *Q*(*s*) are 0, which implies that any non-zero terms *Q*_*ij*_(*s*) are A) strictly proper transfer functions and thus causal, B) descriptions of causal interactions among measured *Y*_*i*_ and *Y*_*j*_. This also means that *Q*(*s*), defined in this way, is unique and has *p* less parameters to identify on its diagonal. This construction of *Q*(*s*) and *P* (*s*) ultimately ensures identifiability [62] under reasonable assumptions of independent input perturbation [39], [40], [24], [41], [42], [43], [44], [45]. Further, if *Q*(*s*) = *W/s*, then we would face two simultaneous challenges in estimation 1) disentangling autoregulatory dynamics (*Y*_*i*_ to *Y*_*i*_) from pairwise interactions (*Y*_*i*_ to *Y*_*j*_), 2) too many unknown parameters in both *Q*(*s*) and *P* (*s*). Lastly, we find that studying the pairwise interactions *Q*_*Ij*_(*s*) can already elicit important functional information about a genetic network, as illustrated by the next two examples.

#### 1) Example: The DSF of an Idealized Incoherent Feed-forward Loop

Consider the following synthetic biology design problem: design and implement an incoherent feed-forward loop. Specifically, we consider implementing a feed-forward loop using the synthetic parts pLac-LasR-CFP-LVA, pLas-TetR-YFP-LVA, and pLas-Tet-RFP-LVA and IPTG, C_3_O_6_H_12_ HSL, and aTc as inputs (see Figure 1). We model the protein concentration of LasR-CFP, TetR-YFP, and RFP as *x*_1_, *x*_2_, and *x*_3_, respectively. We denote the corresponding mRNA species for each of these proteins as *m*_1_, *m*_2_, and *m*_3_. A simple model without any loading effects, describing the dynamics of these states can be written as:

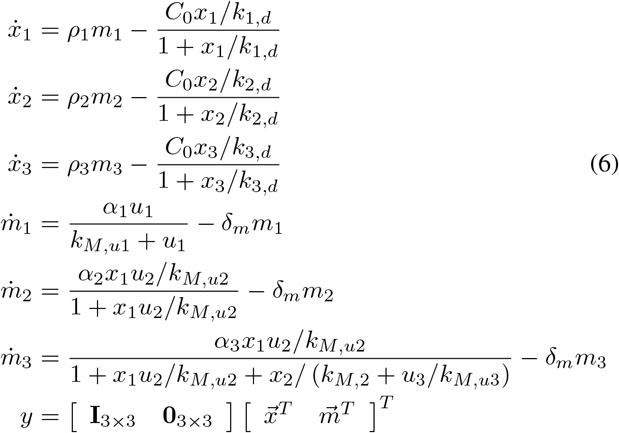

**Fig. 1.**
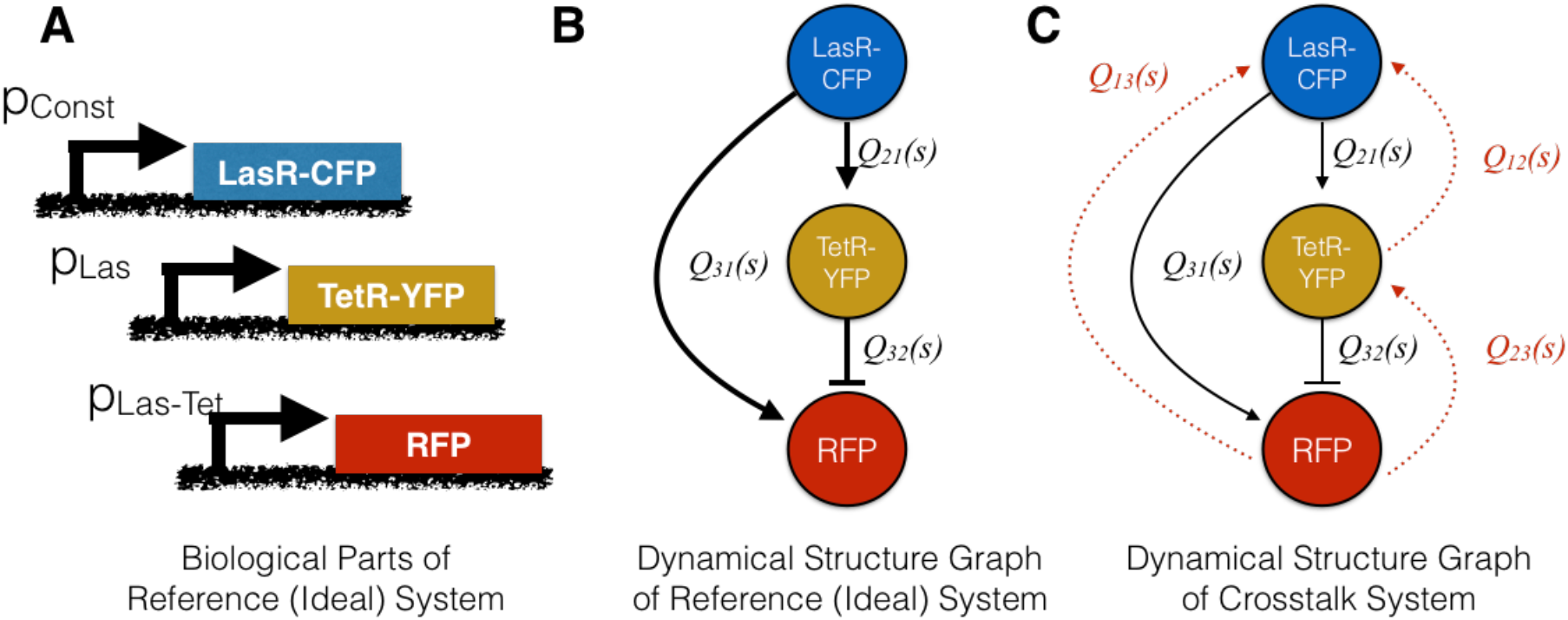
Dynamical structure functions can be used to analyze synthetic gene networks: **(A)** Synthetic biological parts for a incoherent feedforward loop (IFFL) using the LasR activator, the TetR repressor, and reporter proteins CFP, YFP, and RFP. **(B-C)** The dynamical structure graphs of the crosstalk-free IFFL from system (6), in **(B)** and the crosstalk-impacted IFFL from system (7), in **(C)**. Nodes represent measured biochemical species, with black edges denoting causal dependencies stemming from designed interactions, and red edges denoting causal dependencies arising from crosstalk or loading effects. Notice that the dynamical structure captures network models interactions that are not described by the system transfer function *G*(*s*).

The dynamical structure function for this system is derived by taking Laplace transforms and eliminating the hidden mRNA states of *x*_1_, *x*_2_, and *x*_3_, namely *m*_1_, *m*_2_, *m*_3_, see [62] or [66] for a detailed derivation of dynamical structure functions. The network and control structure matrix transfer functions are written (*Q*^*a*^(*s*), *P*^*a*^(*s*)) where *Q*^*a*^(*s*) is written as

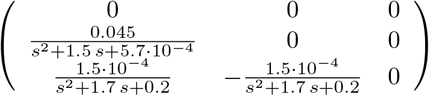

and *P*^*a*^(*s*) is

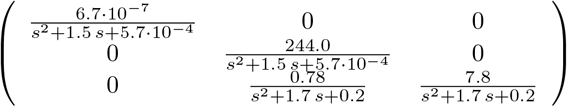

The network, with edge weight functions corresponding to the entries of *Q*^*a*^(*s*), is drawn in Figure 1B. Notice that if we take *s* ∈ ℝ_*>*0_, the sign of the entries in *Q*^*a*^(*s*) coincides with the form of transcriptional regulation implemented by TetR and LasR, respectively. In [67] it was shown that the sign definite properties of entries in *Q*(ℝ_*>*0_) are useful for reasoning about the monotonicity of interactions between measured outputs and how fundamental limits in system performance relate to network structure.

Let us now consider the inverse Laplace transform of *ℒ*^−1^ (*Q*^*a*^(*s*)), we remark that 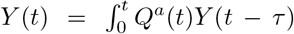 follows from the equation

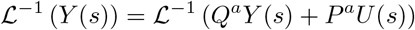

whenever *u*(*t*) ≡ 0 such that *U* (*s*) is 0. This argument holds in general for any system of the form (2). In particular, the entries *Q*^*a*^(*t*) act as convolution kernels, and taken with the integral, define an operator for mapping *y*_*j*_(*t*) to *y*_*i*_(*t*). Most interestingly, we can see that the network structure of this incoherent feedforward loop is *dynamical*, hence our usage of the term *dynamical structure* function to describe the network structure among the measured chemical species *y*(*t*). In this particular case, the time-domain analogue of the dynamical structure (or dynamical structure convolution kernel) is given as

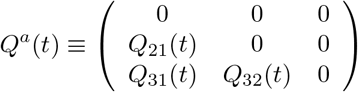

where

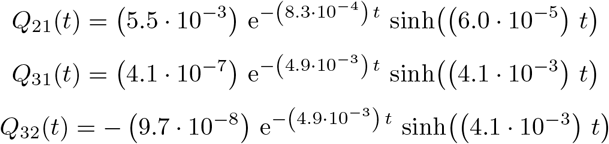

A visualization of each of these impulse kernel functions and their corresponding location in the dynamic adjacency matrix, defined by *Q*^*a*^(*t*), is given in Figure 2. Notice how the activating or repressing nature of genetic regulation is encoded by the positive or negative sign of the corresponding kernel response. In addition to uncovering the Boolean network of interactions between biological states, the dynamical network convolution kernel *Q*^*a*^(*t*) reveals the time-scales of response of each network edge, as well as the amplitude and the rate of decay of the gain from the time of impulse. Similar response profiles can be generated for step function inputs, though finite impulse inputs are typically more common in biological networks. Interestingly, the transfer function *G*^*a*^(*s*) of the system is likewise lower triangular, reflecting the feedforward network topology in the genetic circuit. Specifically, *G*^*a*^(*s*) has a sparsity structure of the form

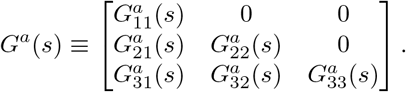

**Fig. 2.**
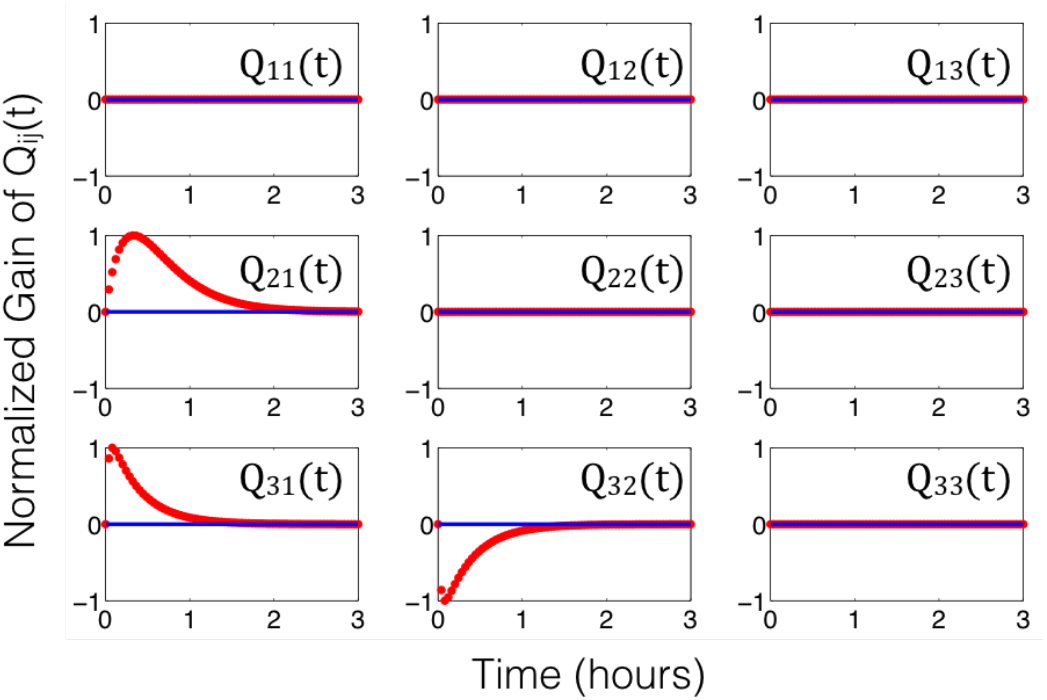
Dynamical structure functions describe how network structure evolves over time (and as a function of frequency): The time-lapse response of the dynamical structure convolution kernel *Q*^*a*^(*t*) = ℒ^−1^ (*Q*^*a*^(*s*)) for the incoherent feedforward loop in system (6). By examining the functional response of each entry in *Q*^*a*^(*t*) (or *Q*^*a*^(*s*)), we see that the network structure of the incoherent feedforward loop in Example II-A1 is a time-evolving, or *dynamic*, entity.

#### 2) Example: The DSF of an Incoherent Feedforward Loop with crosstalk

In prototyping a feedforward loop, it is important to anticipate *in vivo* context effects. We consider the same biocircuit as described in Example II-A1, except now we specifically consider loading effects frequently neglected in the design process of synthetic biology. First, we note that each gene may be susceptible to loading effects [7]. For each gene in Figure 1A, a degradation tag is added, to provide tunability, to the rate of degradation of the protein. Inside the cell, a protease called ClpXP targets these degradation tags and degrades the associated protein. Different tags can be incorporated to modulate the gain of the degradation process. Further, these degradation tags can be subject to mutagenesis experiments, as a means to modulate tunability.

Tunability of degradation introduces a tradeoff in performance. Since the ClpXP protease is a housekeeping protein expressed to form a common pool of proteases for all genes in the cell, there is a limit to the supply of free ClpXP protein in any instant of the cell’s growth cycle. When there are too many degradation-tagged proteins [68], the overloading of the protein degradation queue can trigger unwanted effects such as stress response. More directly, the competition for scarce proteases can induce coupled dynamics or a *virtual* or *indirect* interaction between two genes competing for the same protease pool. Even if the genes were engineered to have no direct transcriptional or translational cross-regulation, the competition for the same protease effectively couples the protein states of both genes. Modifying the above model to account for these type of loading effects yields:

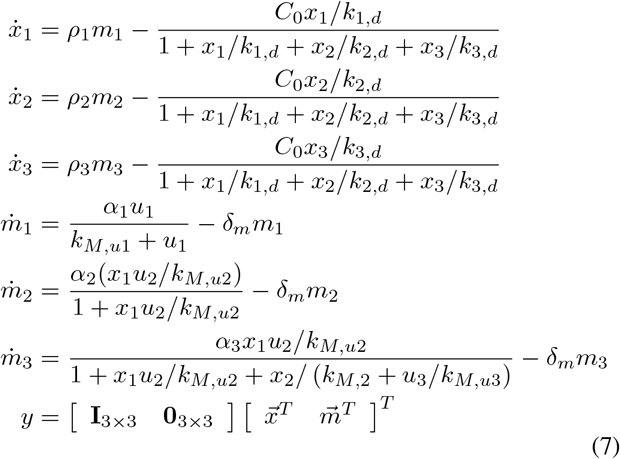

Computing the dynamical structure function, we obtain *Q*^*c*^(*s*)

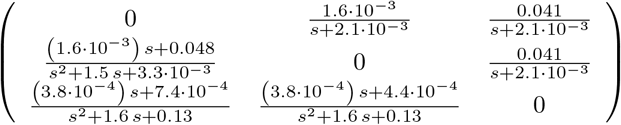

and *P*^*c*^(*s*)

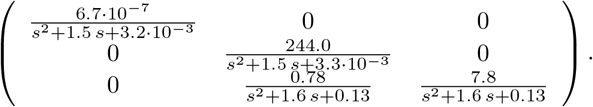

Notice that *Q*^*c*^(*s*) is no longer lower-triangular, but fully connected. Introducing loading effects creates additional coupling between nodes in the network. If the coupling is significant, the *designed* network interactions of the incoherent feedforward loop are overcome by the *crosstalk* network interactions [67], [56], [69], [70], [20], [71], [8]. Thus, the coupling that is introduced into the biochemical reaction network by loading effects is reflected in the structure of (*Q*^*c*^, *P*^*c*^)(*s*).

In contrast, the transfer function of the crosstalk system only characterizes how system outputs causally depend on inputs. In particular, *G*_*c*_(*s*) is also a full matrix like *Q*^*c*^(*s*) of 6th order SISO transfer functions

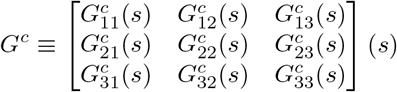

but all structural information about how loading effects cause interference *among* system states is mixed with the information about how outputs causally depend on inputs in *G*(*s*). An identification algorithm of entries in *G*(*s*) will thus be *unable* to quantify the size of crosstalk or interference among system states.

To what extent can the entries of (*Q*(*s*), *P* (*s*)) can be used to quantify the size of crosstalk in a synthetic gene networks? The following theorem shows that the dynamical structure function can be used to quantify crosstalk in biochemical reaction networks.

##### Theorem 1

Let ℒ denote the two-sided Laplace operator. Suppose the states *x*^*c*^ and *x*^*a*^ of the systems (15) are (16) are shifted, so that the origin is a locally asymptotically stable equilibrium point and *Q*^*c*^ and *Q*^*a*^ are the respective dynamical structure functions calculated for each linearized system about the origin. Then

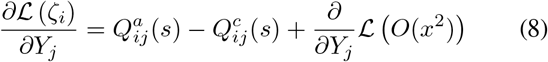

and in particular, if

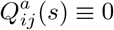

then

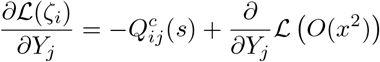

and can be estimated from input output data (*Y* (*s*), *U* (*s*)). The proof of this theorem is provided in the Supplementary Information. There are two ways to apply this result. First, if one has an idealized or reference dynamical structure model of the circuit, then this can be compared to the dynamical structure model estimated from the data. Second, in the absence of a reference model, the dynamical structure model can be estimated directly from data to observe new edge functions or gain mismatch directly from the data-driven dynamical structure model. In the latter scenario, if the crosstalk interactions are mediated on an existing edge, it will not be possible to separate the magnitude of the crosstalk from the existing (intended) dynamics of a given active edge in the network. However, the discovered edge dynamics can be compared to the intended behavior, e.g., intentional activation or repression over a certain growth phase of the cells.

## III. Direct Estimation Algorithm for Dynamical Structure Functions

In this section, we will introduce a direct estimation algorithm for estimating the dynamical structure function. The dynamical structure function is a tuple of matrix transfer functions (*Q*(*s*), *P* (*s*)) and can be directly estimated from experimental or simulation data so long as the data and the conditions of the experiment satisfy the assumptions of the following theorem, proved in [62].

### Theorem 2

Given a *p* ×*m* transfer function *G*(*s*), dynamical structure reconstruction is possible from partial structure information if and only if *p* − 1 elements in each column of [*Q*(*s*) *P* (*s*)]^*\**^ are known that uniquely specify the component of (*Q, P*) in the nullspace of [*G*(*s*)^*\**^ *I*].

It follows from this theorem that without additional structural information about the columns of the matrix [*Q*(*s*) *P* (*s*)]^*\**^, it is not possible to identify (*Q*(*s*), *P* (*s*)). In synthetic gene circuits with complex internal network interactions encoded by *Q*(*s*), we can still solve for the structure and parameters of *Q*(*s*), as long as enough elements of *P* (*s*) are known. For example, targeted gene knockdowns (CRISPRi) or knockouts (engineered genomic mutations) can be engineered so that *P* (*s*) is a diagonal matrix transfer function. That is, *all genes or biological states in the network of interest* are A) monitored by some measurement channel over time and B) independently perturbed by a diagonal element in *P* (*s*). These are necessary and sufficient conditions for reconstruction of *Q*(*s*) and *P* (*s*) and align with the conditions developed by Sontag, Kholodenko, Wolkenhauer, Kang, Prabakaran et al. for network reconstruction of full state measurement systems [39], [40], [24], [41], [42], [43], [44], [45].

Under the above premises, the task is to estimate the diagonal transfer function entries of *P* (*s*) and to estimate all off-diagonal entries of *Q*(*s*). Recall from the derivation in Section II-A that the diagonal entries of *Q*(*s*) are set to 0 by subtracting the diagonal entries of the precursor *W* (*s*) transfer function matrix from *W* (*s*). Further, by left-multiplying (*sI* − *D*)^−1^ with *W* (*s*) − *D*(*s*), this guarantees that the representation for *Y* = *QY* +*PU* yields a unique *Q*(*s*) and *P* (*s*). The entries of *Q*(*s*) and *P* (*s*) are all strictly proper rational transfer functions and thus encode causal dynamics.

There are two approaches to solve for *Q*(*s*) and *P* (*s*). The first is to estimate the transfer function *G*(*s*) using a standard transfer function estimation routine, followed by inversion of *G* and calculation of the entries of *P*_*ii*_(*s*) and subsequently the entries of *Q*(*s*). This approach has the drawback of relying on inversion of the matrix transfer function *G*(*s*), which often requires symbolic inversion and is thus prone to numerical instability and scaling issues for larger networks.

The second approach, which we propose here and develop code for, is to identify the dynamical structure function directly from data by writing the model estimation problem in normal form. First, we can estimate the discrete time approximations of *Q*(*s*) and *P* (*s*), as 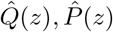, which will follow the same identifiability conditions [62].

Specifically, we have that

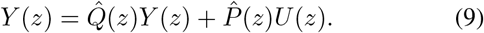

which given that 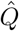 and 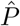 share the same denominator, we can multiply the characteristic polynomial of 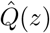 on both sides, to obtain

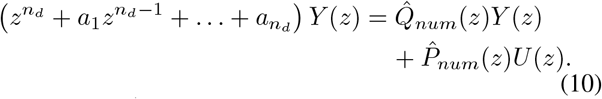

The matrices 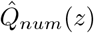 contain unknown coefficients like 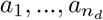 and so we can express the known quantity 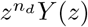 after inverse *Z*-transforming to obtain

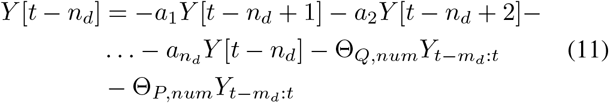

which can be written in normal form as

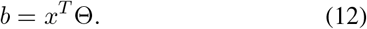

This equation can be stacked for multiple time traces, collected from different conditions or experimental replicates, or by staggering the time horizon, to obtain the stacked normal form equations

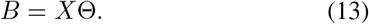

where *B* and *X* are known values of time-series data and Θ contains the coefficients determining the characteristic polynomial and the numerators of the dynamical structure function. Once these estimates are obtained, a standard discrete-to continuous transformation can be used to estimate *Q*(*s*) and *P* (*s*). The steps of our algorithm are summarized in Algorithm 1 and the full MATLAB code is provided at the Github repository https://github.com/YeungRepo/NetRecon.

A couple remarks are in order. First, there are two distinct hyperparameters that require optimization in generating the estimates for *Q*(*s*) and *P* (*s*): 1) *n*_*d*_, the order of the characteristic polynomial and 2) the selection of the number of subsampled timepoints *h*_*max*_ in a given time-series traces. The optimal value of *h*_*max*_ will depend on the dataset, to ensure the condition number of the matrix *X* must be small. In high resolution time-series measurements, a small or short subsampled horizon *h*_*max*_ may produce virtually identical data if the transient has a slow rate of change, which can result in ill-conditioning of *X*. We optimize *n*_*d*_, *h*_*max*_ to minimize the n-step L1 prediction error across all *N*_*exp*_ experimental samples. In practice, we observe that this criteria, by necessity, guarantees an appropriate selection of *h*_*max*_ as well as a lower optimal *n*_*d*_ value.

Secondly, the routine described above may appear to be easily cast as a classic transfer function estimation routine, similar to the tools developed by [73] in MATLAB. The primary difference in this procedure from a standard transfer function estimation procedure for

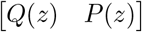

is that we must impose structural information about *Q*(*z*) and *P* (*z*) on the estimation process. This results in a structured system identification problem, which typically is non-convex and difficult to initialize properly. Specifically, the diagonal entries of the estimated *Q*(*z*) and the off-diagonal entries of *P* (*z*) must be exactly 0, as per the premises of [62]. Again, these structural constraints guarantee a unique representation of the dynamical structure function and identifiability of the model. When using standard transfer function estimation tools (in MATLAB’s System Identification toolbox), we found repeatedly over thousands of numerical trials that imposing structural constraints as model priors resulted in A) models with extremely poor n-step prediction capacity (forecasting) or B) models with extremely poor 1-step fit scores. This motivated the development of a direct estimation algorithm, that mirrors the standard estimation of an discrete-time transfer function model, but where *structural constraints* of *Q*(*z*) and *P* (*z*) are directly encoded into the formulation of the normal form of equations (Lines 19-28, 32 and 36 of Algorithm 1).

### Algorithm 1: Direct QP Estimation Algorithm

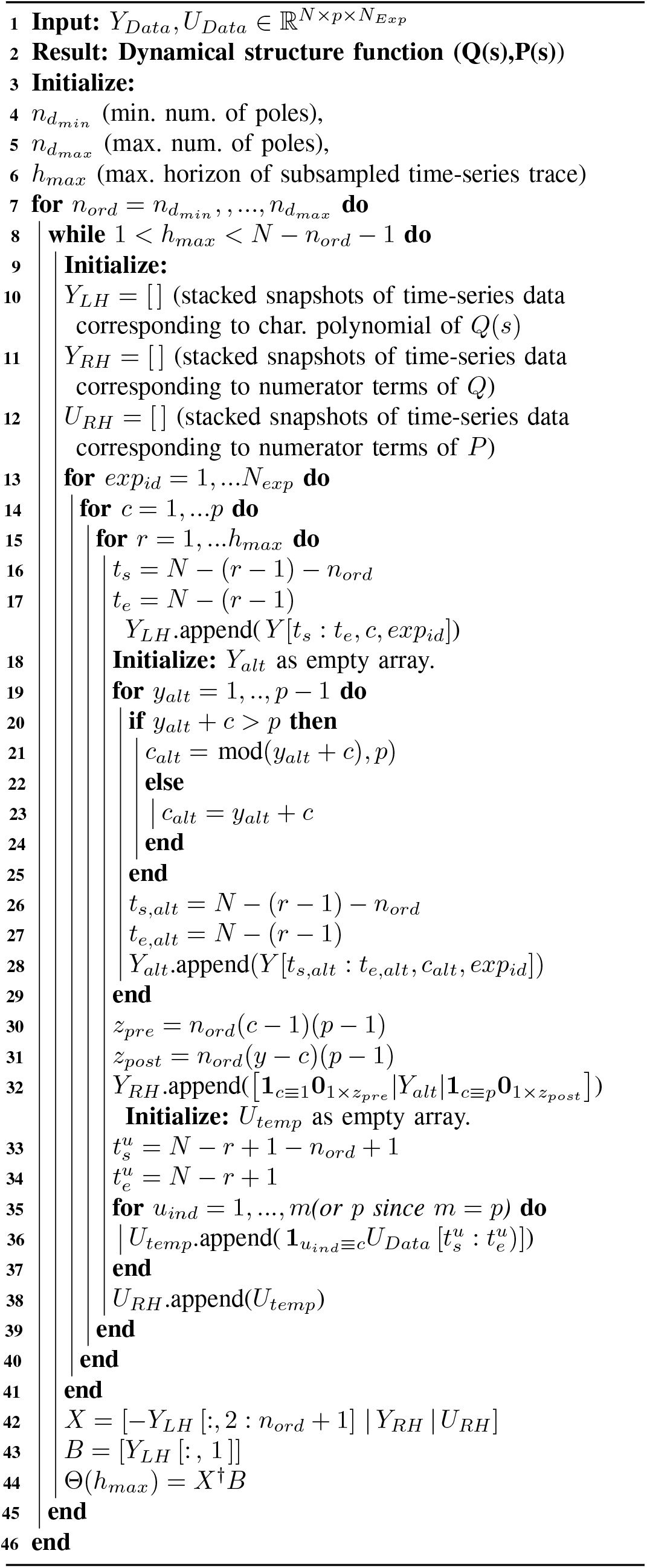

## IV. The Dynamical Structure of an *in vitro* Genelet Repressilator

We now illustrate the process of data-driven estimation of dynamical structure functions, using experimental data. In this section, we take as a first test case the synthetic genelet repressilator developed by Kim and Winfree [72]. The genelet repressilator consists of three DNA switches that repress one another through indirect sequestration. Specifically, each DNA switch transcribes its mRNA product only when its activator strand binds to complete its T7 RNA polymerase promoter sequence. The RNA product produced from each DNA switch, in turn, acts as an inhibitor to the downstream switch by binding to the downstream switch’s DNA activator molecule. Thus, by sequestering the DNA activator from completing the T7 RNA polymerase promoter region, the mRNA product of the upstream switch inhibits activation of the downstream switch. Figure 4A shows the mechanistic design of the genelet switch.

**Fig. 3.**
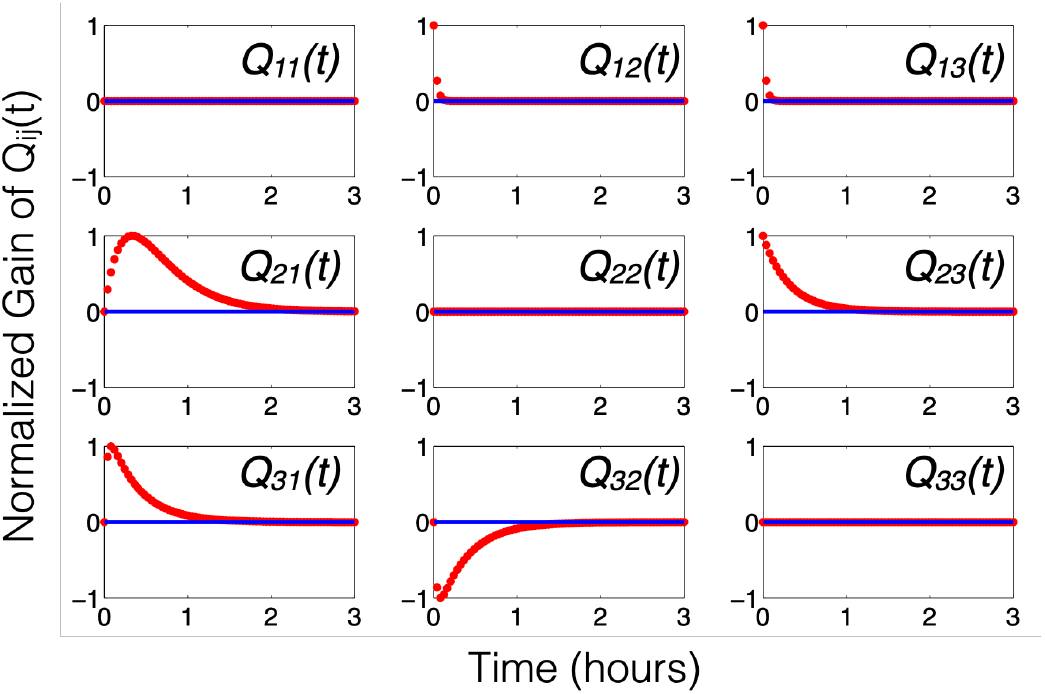
Dynamical structure functions describe how network structure evolves over time (and as a function of frequency): The time-lapse response of the dynamical structure convolution kernel *Q*^*c*^(*t*) = ℒ^−1^ (*Q*^*c*^(*s*)) for the incoherent feedforward loop in system (7). By examining the functional response of each entry in *Q*^*c*^(*t*) (or *Q*^*c*^(*s*)), we see that the network structure of the incoherent feedforward loop in Example II-A1 is a time-evolving, or *dynamic*, entity.

**Fig. 4.**
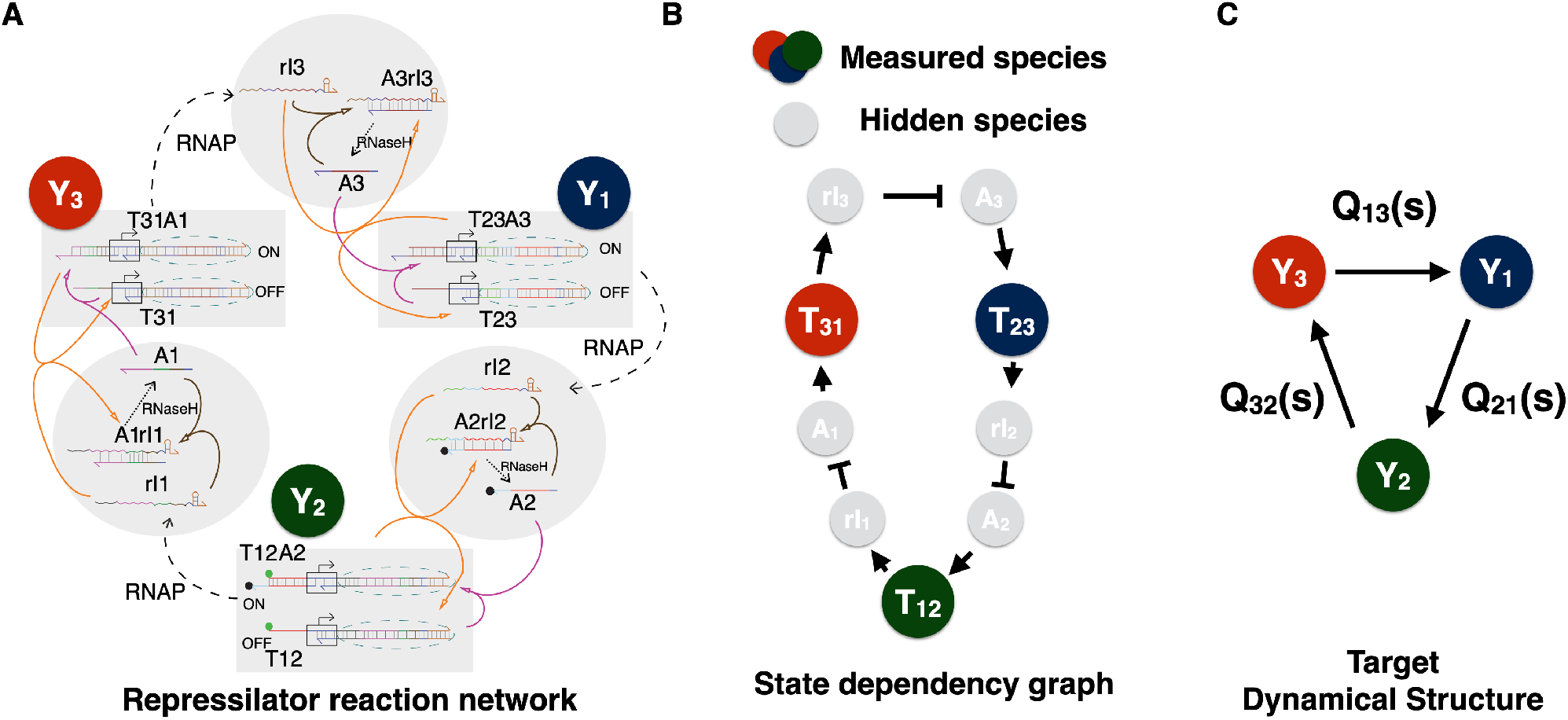
Network representations of a synthetic genelet repressilator: **(A)** A reaction network using push-arrow reaction notation of the synthetic genelet repressilator. **(B)** A diagram representing the reaction dynamics in panel **(A)** as state dependencies from a nonlinear ODE model in [72]. **(C)** The dynamical structure of the repressilator (without inputs), with nodes representing measured chemical species and edge weights corresponding to entries in *Q*(*s*).

The genelet switch relies heavily on RNase H to degrade any activator-mRNA inhibitor complexes. Without degradation, the binding of activator to mRNA inhibitor is much faster than unbinding and so sequestration is effectively irreversible. Thus, in order for the repressilator to function properly, RNase H must degrade its target substrates sufficiently fast. If RNase H is saturated with high levels of a particular substrate, this slows the degradation of other substrates, creating a crosstalk interaction between competing DNA-RNA complexes.

By performing network reconstruction on the genelet repressilator, we can determine how much crosstalk exists in the biocircuit. Further, we can validate our dynamical structure reconstruction algorithm in an *in vitro* setting, by *deliberately attenuating one of the components to create a gain imbalance*. We can see if the reconstruction process recovers the deliberate imbalance we introduce into the genelet repressilator, even when simply measuring local perturbations of an operating point for a normally oscillatory circuit.

To reconstruct *Q*(*s*) and *P* (*s*), we performed a single experiment with three perturbations applied in series [39], [40], [24], [41], [42], [43], [44], [45], [62]. To perturb each switch we pipetted a small perturbative concentration of DNA inhibitor (a DNA analogue of RNA inhibitor). Since DNA is not degradable in a T7 expression system by RNase H, it effectively acts as a step input since it binds to DNA activator and does not degrade. In this way, our perturbation design ensures sufficiency of excitation and independent perturbation of each activator (and downstream switch), thereby satisfying the identifiability conditions in [62] and the persistence of excitation conditions described in [73]. Further, we attenuated the concentration of the third switch *T*_31_ by 20%, to *create a deliberate gain imbalance for evaluating our reconstruction algorithm*.

A detailed model of the repressilator can be found in the supplement of [72]. Since the derivation is lengthy, it suffices to write the idealized dynamical structure function *Q*^*a*^(*s*) of this system, corresponding to the detailed model provided in Supplementary Section 1.6 [72]. The structure is obtained by linearizing the system, transforming into the Laplace domain, eliminating hidden variables to obtain the following:

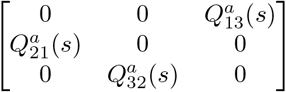

reflecting the cyclic structure of the system. This represents an idealized model of the system. As stated in Theorem **??** every entry where 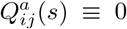, the corresponding entry in *Q*(*s*) estimated directly from experimental data, will be a crosstalk interaction present in the network. Here *Q*(*s*) is used to denote *Q*^*c*^(*s*), the dynamical structure function estimated directly from data.

The experimental data used to fit *Q*(*s*) and *P* (*s*) are plotted in Figure 5, along with their respective fits. For each row *i* of *Q*(*s*), we use *Y*_*j*_, *j* ≠ *i* and *U*_*i*_ as inputs and *Y*_*i*_ as the output for a direct MIMO *p* ×1 transfer function estimation problem. The impulse response for the convolution kernel *Q*(*t*) of the reconstructed *Q*(*s*) is plotted in Figure 6.

**Fig. 5.**
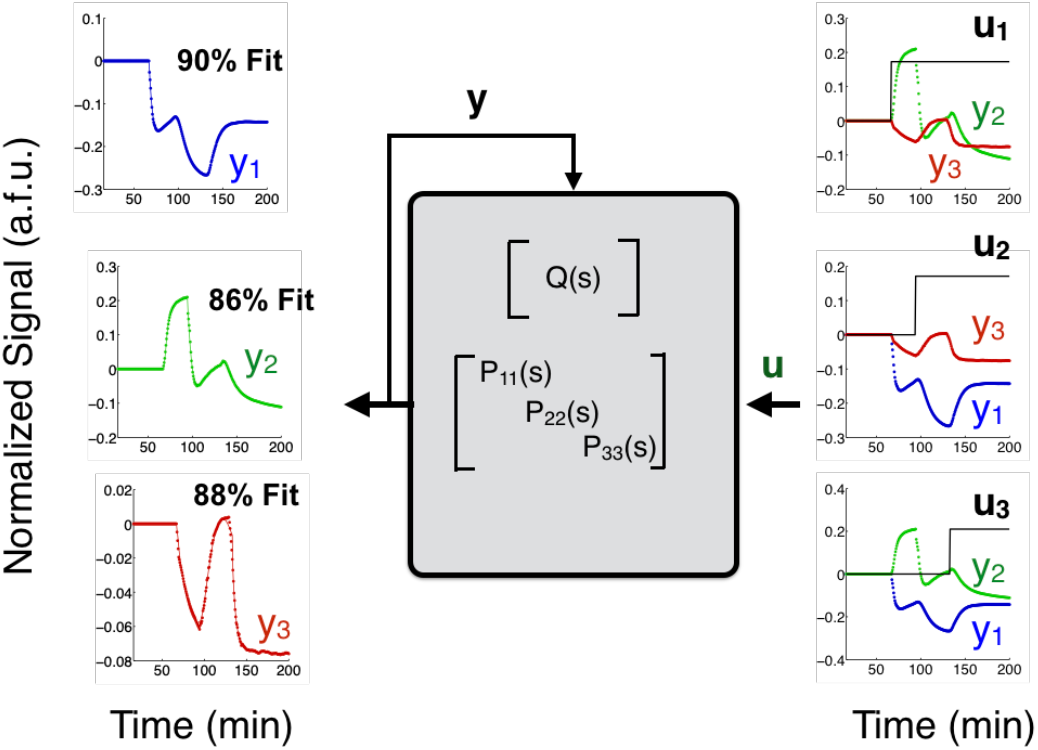
Time-series experimental data from *in vitro* network perturbation experiments of a T7 RNAP genelet repressilator. Three outputs are measured simultaneously, *y*_1_, *y*_2_, and *y*_3_, corresponding to DNA switches *T*_31_, *T*_12_ and *T*_23_. DNA homologues of the RNA inhibitors *rI*_*j*_ *j* = 1, 2, 3 are injected at small concentrations to provide a step input perturbation to the corresponding component *Y*_*j*_ in the genelet circuit.

**Fig. 6.**
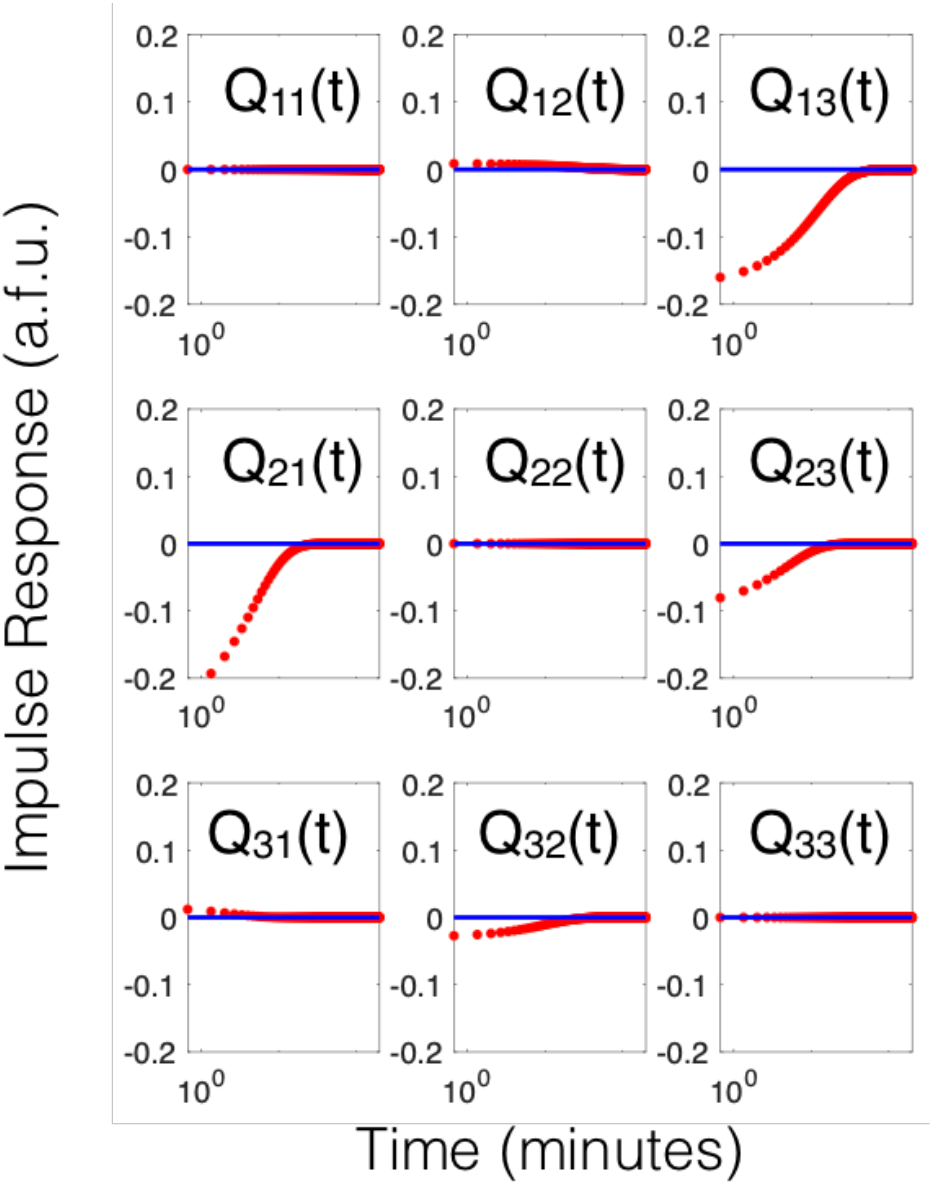
Impulse response of the estimated convolution kernel *Q*(*t*) matrix. *Q*(*s*) is estimated directly from experimental data, transformed into the frequency domain, and simulated in time for *t* = 0 to *t* = 300 minutes.

If we compute the corresponding ℋ_*∞*_ gain of each entry in *Q*_*ij*_(*s*) and scale by the maximum gain, we obtain

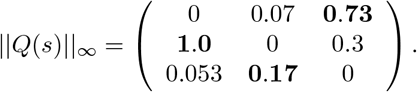

We see significant crosstalk on the edge *Q*_23_(*s*) and minor crosstalk from entries *Q*_31_(*s*) and *Q*_12_(*s*). This crosstalk need not occur simultaneously, since the ℋ_*∞*_ gain calculates the worst-case or maximum gain over all possible frequencies. With the exception of *Q*_23_(*s*), all other crosstalk entries have strictly smaller ℋ_*∞*_ gain than the designed edge. Examining the impulse response of the convolution kernel confirms these observations; the crosstalk edge *Q*_23_(*t*) has a larger impulse response than *designed* edge *Q*_32_(*t*).

As intended in the design of the experiment, our estimated network model shows a gain imbalance between the *designed* edges *Q*_32_(*s*), *Q*_13_(*s*) and *Q*_21_(*s*). In order for the oscillator to perform properly, it needs to have approximately the same gain along each edge in the network. This example verifies that our reconstruction algorithm can identify important functional dynamics of a genetic circuit; especially for debugging purposes. The linearization scheme is valid, so long as we model fluctuations in dynamics from a nominal initial condition, even if the initial condition is not stable or leads to oscillatory dynamics Our results here illustrate how linearized models can provide insight into local dynamics. In this simple, controlled dataset, we know we can increase the gain of the edge in *Q*_32_(*s*) by adjusting the binding affinity of the activator DNA with its inhibitor RNA, or by increasing the concentration of the corresponding downstream switch *T*_31_. Notice that this design insight may not be obvious by direct examination of experimental trajectories of each switch in Figure 5. As long as we have an idealized network model, we can measure the deviation from that model *Q*^*a*^(*s*) in the network model identified from the data *Q*^*c*^ = *Q*(*s*) and identify edges or nodes in our network that need tuning.

## V. The Dynamical Structure of an *in vivo* Transcriptional Event Detector

We now introduce a new transcriptional event detector circuit, one that is designed, built, and constructed for illustrating the use of our dynamical structure estimation algorithm in *in vivo* circuit design. Event detectors are useful because of their ability to perform temporal logic. Making temporal logic decisions enable applications such as programmed differentiation, where the goal is to perform some operation based on a combinatorial and temporal sequences of events that dictate cell fate.

So far there are two demonstrations of temporal logic gates: 1) a temporal logic gate that differentiates start times of two chemical outputs [76] and 2) a molecular counter that counts the number of sequential pulses of inducers [77]. Both event detectors use serine integrases to perform irreversible recombination, while [77] demonstrates the use of transcription-based event detecting to perform event counting. The advantage of an integrase-based approach is the persistent nature of DNA-based memory. At the same time, the drawback of integrase-based event detection is that it is limited to one-time use.

In contrast, transcription based event detectors use proteins instead of DNA to encode a memory state [77], [78]. The advantage of a transcription-based event detector is that proteins are labile, since they are diluted through cell growth or can be tagged for degradation. Thus, a transcriptional event detector’s memory state can be reset after some period of time. On the other hand, maintaining protein state over multiple generations is metabolically expensive [71] and the dynamics of the circuit can become sensitive to production and growth phase of the cells. Therefore, a transcription based event detector biocircuit must be designed with precise timing, balance of production rates, and carefully tuned gain of each transcriptional regulator. This presents a suitable application for our network reconstruction algorithm.

### A. Designing a transcriptional event detector

We designed our transcriptional event detector to be made of two constitutively expressed relay genes, AraC and LasR, and an internal toggle switch. The two relay genes transmit the arrival of two distinct induction events (arabinose and HSL) to relay output promoters pBAD and pLas respectively, which drive production of a fluorescent response in two relay promoters. To record these induction events historically, the output of each relay gene is coupled to one of two combinatorial promoters (pBAD-Lac or pLas-Tet) in a toggle switch. Each combinatorial promoter implements NIMPLY logic, e.g. pBAD-Lac (pLas-Tet) expresses TetR (LacI) only when arabinose (HSL) and AraC (LasR) are present and LacI (TetR) is absent. Thus, when one analyte (e.g. arabinose) arrives, it triggers latching of the toggle switch only if the toggle switch is unlatched to begin with or the prior latching protein state has been diluted out. The relay outputs thus transmit the *current or recent* induction event state while the toggle switch maintains the *historical* induction event state. Depending on the order of arrival of each inducer, we obtain different biocircuit states. Figure 7 details the genetic elements in the event detector biocircuit and the designed component interaction network.

**Fig. 7.**
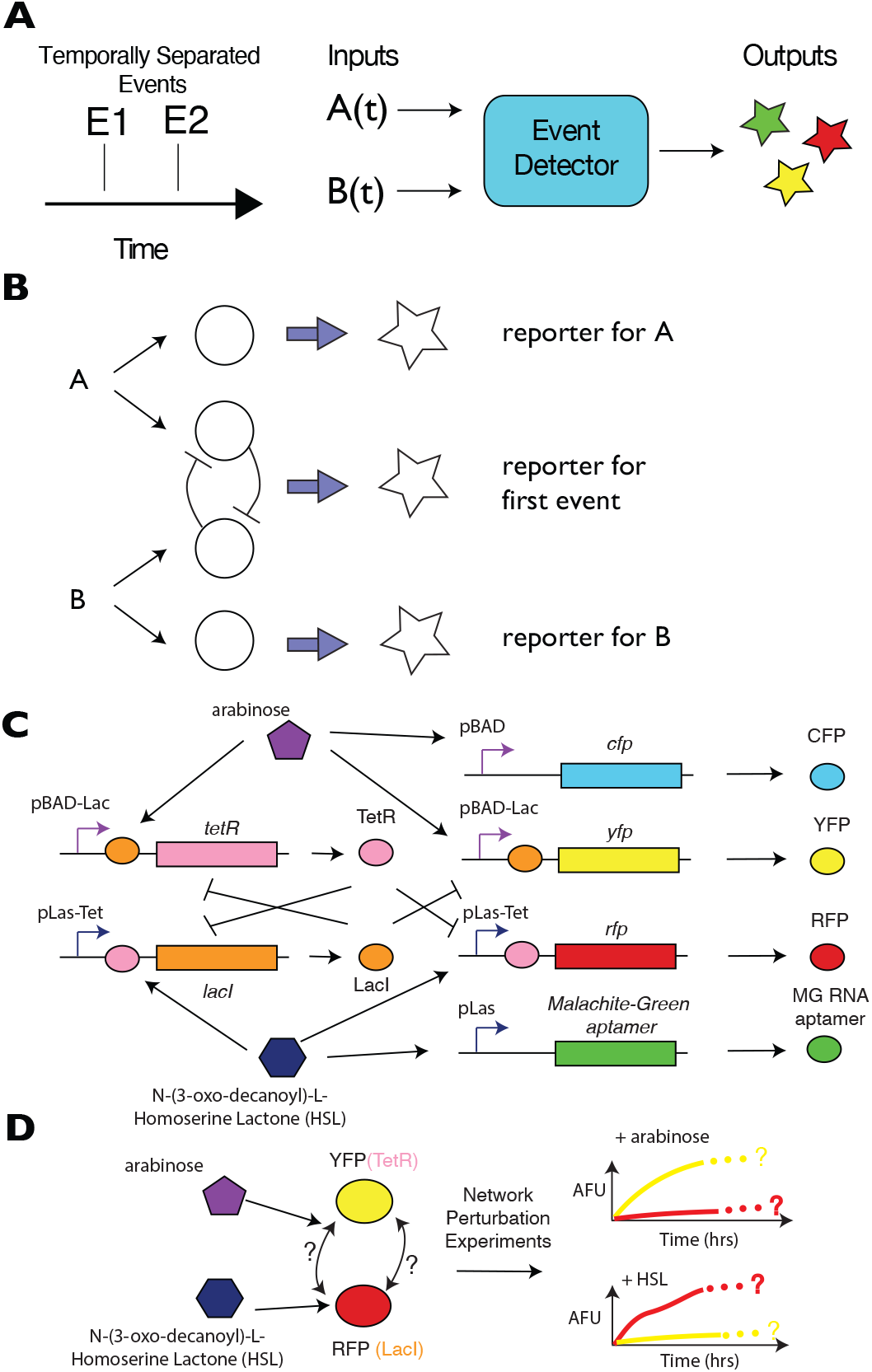
(**A**) Left: We design an event detector to determine the identity and relative ordering of two events *E*_1_ and *E*_2_ occurring within a finite time horizon. (**B**) A schematic showing the logic of the circuit for the event detector. Arrival of event type A triggers transient reporter for A (top) and latching of the toggle in a A-dominant state as a memory state. Similarly, arrival of event type B triggers transient reporter for B (bottom) and latching of the toggle in a B-dominant state as a memory state. (**C**) A diagram showing the synthetic biocircuit parts used to implement the network architecture in (**B**). **D)** The arabinose and HSL inducers independently perturb distinct elements of the memory module in the event detector; a network model of the dynamic graph of the event detector can be reconstructed using dynamical structure function reconstruction experiments.

We can write down an idealized model for the event detector (assuming no crosstalk), assuming first order degradation and production, with Hill functions encoding the NIMPLY logic of each promoter in the memory module.

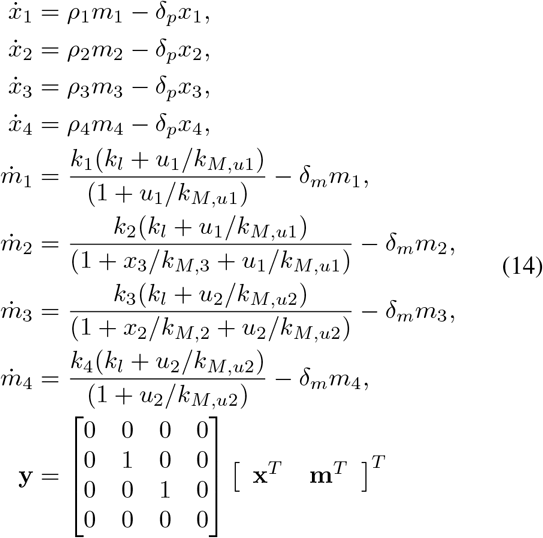

where the measured outputs of the system are *y*_*i*_ = *x*_*i*_, *i* = 2, 3, *ρ*_*i*_ is the translation rate of *m*_*i*_ into *x*_*i*_, *δ*_*p*_ is the effective dilution rate of *x*_*i*_, *i* = 1, …, 4, *δ*_*m*_ is the combined dilution and degradation rate of *m*_*i*_, *i* = 1, …, 4, *k*_*M*_, *u*_*i*_ is the Michaelis constant for *u*_*i*_, *k*_*l*_ is the leaky catalytic transcription rate, *k*_*i*_ is the catalytic transcription rate for *m*_*i*_, and *u*_1_, *u*_2_ are arabinose and HSL, respectively.

Again, the dynamical structure function for this system is calculated by linearizing the system about a nominal initial condition, (*x*_0_, *m*_0_), taking a Laplace transform and solving out the hidden variables *m*_1_, …, *m*_4_. We present a simplified case here, assuming algebraic symmetry of the parameters *k*_*i*_ = *k, ρ*_*i*_ = *ρ, k*_*M,i*_ = *k*_*M*_ as it does not qualitatively change the structure of (*Q*(*s*), *P* (*s*)). We obtain:

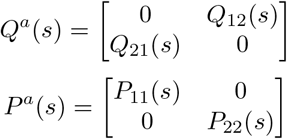

where *P*_*ii*_(*s*)= *ρ/* (*δ* _*m*_ + *s*) (*δ* _*p*_+ *s*) for *i* = 1, 2 and

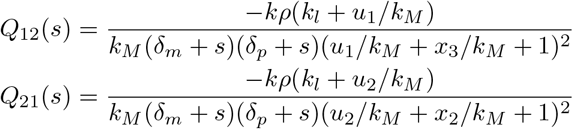

In the absence of protein degradation, *Q*_12_(*s*) and *Q*_21_(*s*) can be approximated with first-order SISO transfer functions. These expressions for *Q*(*s*) and *P* (*s*) are for the idealized dynamical structure function of the *alternative* system. Notice that *Q*_12_(*s*) and *Q*_21_(*s*) are strictly negative transfer functions, indicating the repression present in an idealized simulation of the event detector circuit. This is the intended *dynamical network* structure of the event detector, in the absence of all genetic crosstalk or context effects.

Depending on the abundance of transcription factors such as LacI, TetR, and AraC, as well as commonly shared transcriptional and translational proteins, the *actual* dynamical structure function *Q*^*c*^(*s*) may not exhibit monotonic repression or may even unveil unwanted interactions. We can investigate these interactions under a range of conditions with dynamical structure estimation.

We constructed a biological implementation of the event detector, using the design specified in Figure 7. The logical components containing the relays and the memory module were encoded on to a plasmid vector with a kanamycin resistance marker and a ColE1 (high copy) replication origin. The fluorescent reporter elements with the relay promoters and readouts for the toggle switch were encoded on a plasmid vector with chloramphenicol resistance and the p15 replication origin.

### B. Event Detector Latching Experiments

We evaluated the performance of our transcriptional event detector circuit using a temporal logic test. A standard temporal logic experiment for any two-input event detector is to evaluate the effect of varying the order of presentation of two input signals. In one test, we present the first input, arabinose, for 7.5 hours, followed by induction of the second input, a homo-serine lactone (HSL) quorum sensing molecule to activate the pLas-Tet promoter. In the second test, we swap the order of the inputs, presenting HSL quorum sensing molecule to the event detector for 7.5 hours, then present arabinose inducer as a second input. Both tests evaluate the ability of the memory module of the event detector to latch in the correct state in response to the first input, followed by a challenge to ignore the second input signal while the relays detect and read out the second input signal. The data for both of these *in vivo* tests is plotted in Figure 8B-C.

**Fig. 8.**
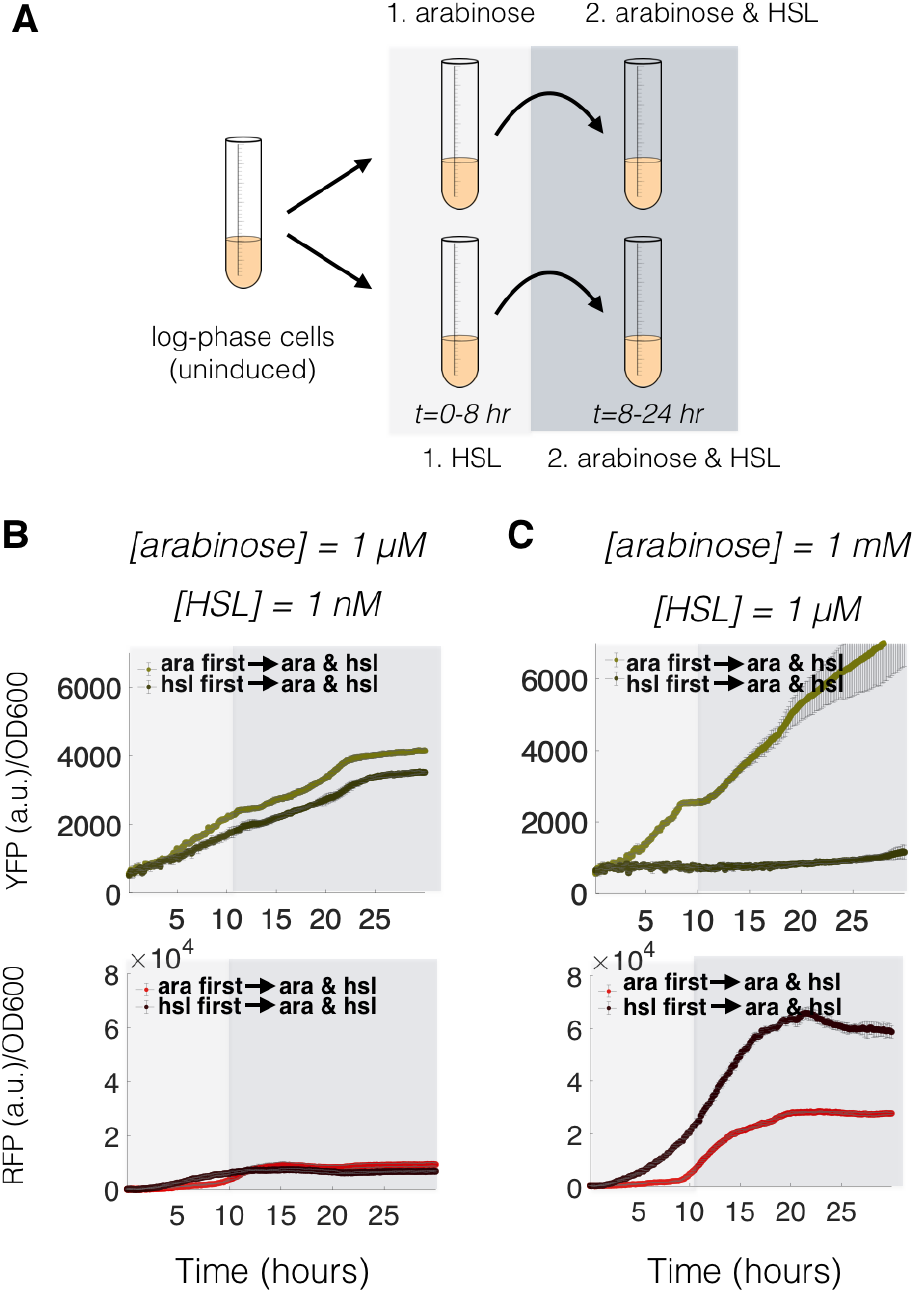
A plot of data from *in vivo* plate reader experiments, testing the temporal logic properties of the event detector diagrammed in Figure 8. Notice that at 1 *µ*M HSL and 1*mM* arabinose, the event detector functions properly, expressing different levels of YFP and RFP depending on the the order of arrival of arabinose and HSL. At 1 nM HSL and 1 *µ*M arabinose induction concentrations, the temporal logic properties of the event detector completely are abolished.

The event detector showed the correct latching response in all tests at standard maximum induction concentrations of arabinose (1 mM) and working induction concentrations of 1 *µ*M HSL. For example, Figure 8C shows that when the event detector is given arabinose followed by HSL, it generates the correct fluorescent response of yellow fluorescent protein, with lower expressions level of RFP. Conversely, when we add HSL first, followed by arabinose, RFP signal ramps up immediately beginning as early as 1-2 hours after induction while YFP expression is abolished to background levels.

We tested a variety of combinations of high and low concentrations for arabinose and HSL. When the concentration of HSL was decreased to 1 nM, we observed consistent leaks in the memory module in either the YFP channel or the RFP channel. Decreasing arabinose down to 1 *µ*M still allows for latching of high YFP expression, but in the presence of 1 *µ*M HSL, any arabinose latching is reversed by HSL induction (data not plotted). Conversely, when we attenuate HSL induction to 1 *nM*, HSL does not prevent arabinose from reversing a HSL latch on the the memory module, see Figure 8B. This leak is significant enough in the 1 nM HSL induction level that the difference in signal between the arabinose-HSL induction scenario versus the HSL-arabinose induction scenario vanished. This temporal logic response profile is evident of a glitch in the event detector circuit that occurs at lower HSL and arabinose concentrations.

### C. Network Reconstruction Experiments to Debug Circuit Failure

We conducted 4 *in vivo* network reconstruction experiments (2 inducers versus 2 concentrations), recording time-series data of the memory module relay elements, YFP and RFP. The memory module is designed using two hybrid promoters, so from a design standpoint, verification of the memory module was most critical. The arabinose inducer targets the pAra-Lac promoter, while the HSL inducer targets the pLas-Tet promoter (see Supplementary Information for sequences).

As shown in the model (14) of the event detector, the actual event detector we constructed exhibits nonlinear response. However, for any one parametric concentration regime, e.g. at a fixed arabinose or HSL concentration, the response of the system behaves similar to that of a linear system. Thus, we estimated a dynamical structure function for both conditions of the reconstruction experiment. The one-step accuracy in fitting dynamical structure models to the low gain condition (1 *µ*M arabinose, 1 nM HSL) and high gain condition (1 mM arabinose and 1 *µ*M HSL) were 99.996% and 99.995% respectively.

As in the case of the genelet repressilator, we can plot a dynamical network graph for the *in vivo* event detector to understand how the memory module components labeled by YFP and RFP, representing TetR and LacI respectively, interact with each other. A movie visualizing the dynamics of the edges of the graph is available for download (see Supplementary Information). Each edge represents the convolution kernel response of the edge to an impulse applied to that input. All responses are superimposed to form a dynamical graph. Snapshots of the graph are plotted in Figure 10, while time-lapse responses of the weights of each edge are plotted in Figure 9. Again as with the repressilator, we can see that the regulatory nature of edges in the event detector’s memory module manifests as two edges with negative or positive values indicating repression or activation, respectively.

**Fig. 9.**
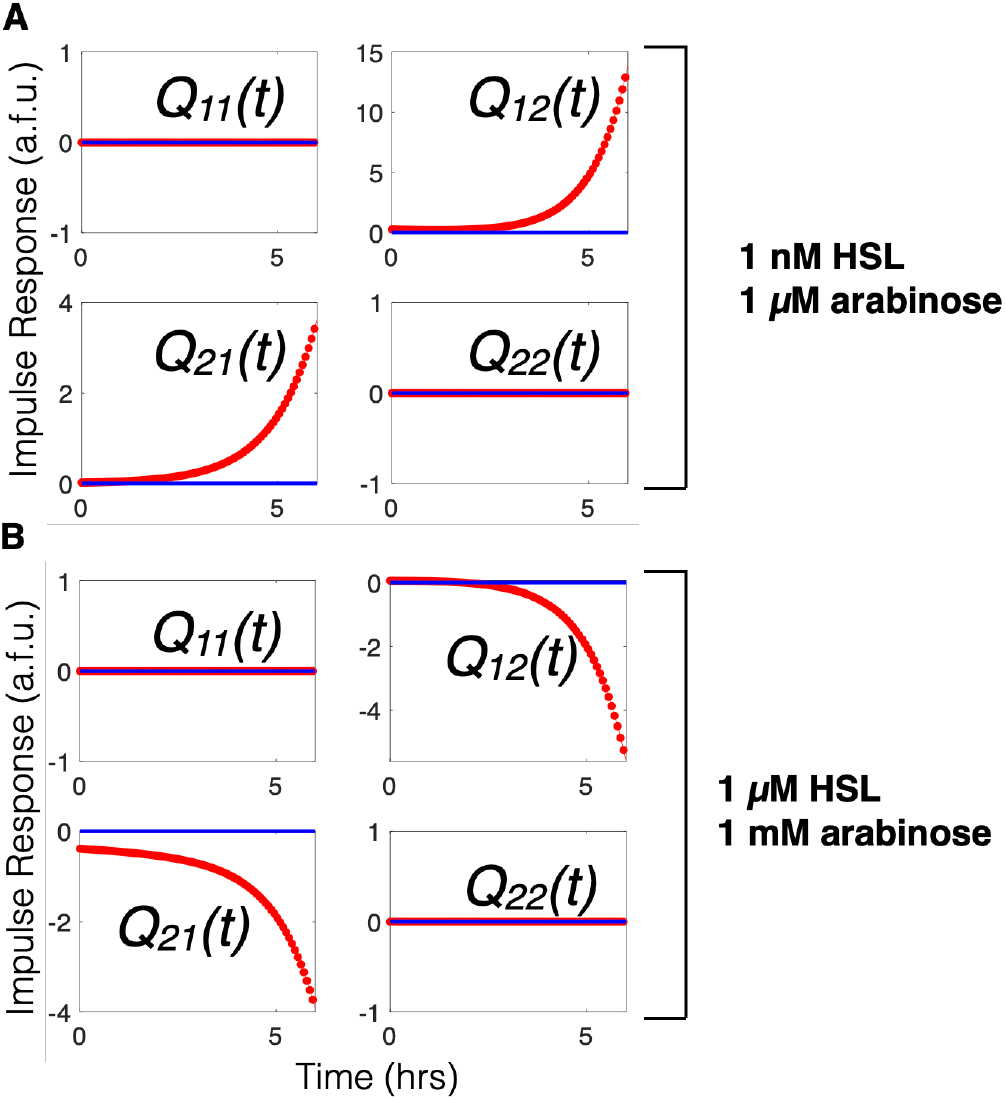
Impulse response of the estimated convolution kernel *Q*(*t*) matrix when the event detector biocircuit is induced with (A) 1 nM HSL and 1 *µ*M arabinose or (B) 1 *µ*M HSL and 1 mM arabinose. *Q*(*s*) is estimated directly from experimental data, transformed into the frequency domain, and simulated in time as a function of hours from arrival time of an inducer input.

**Fig. 10.**
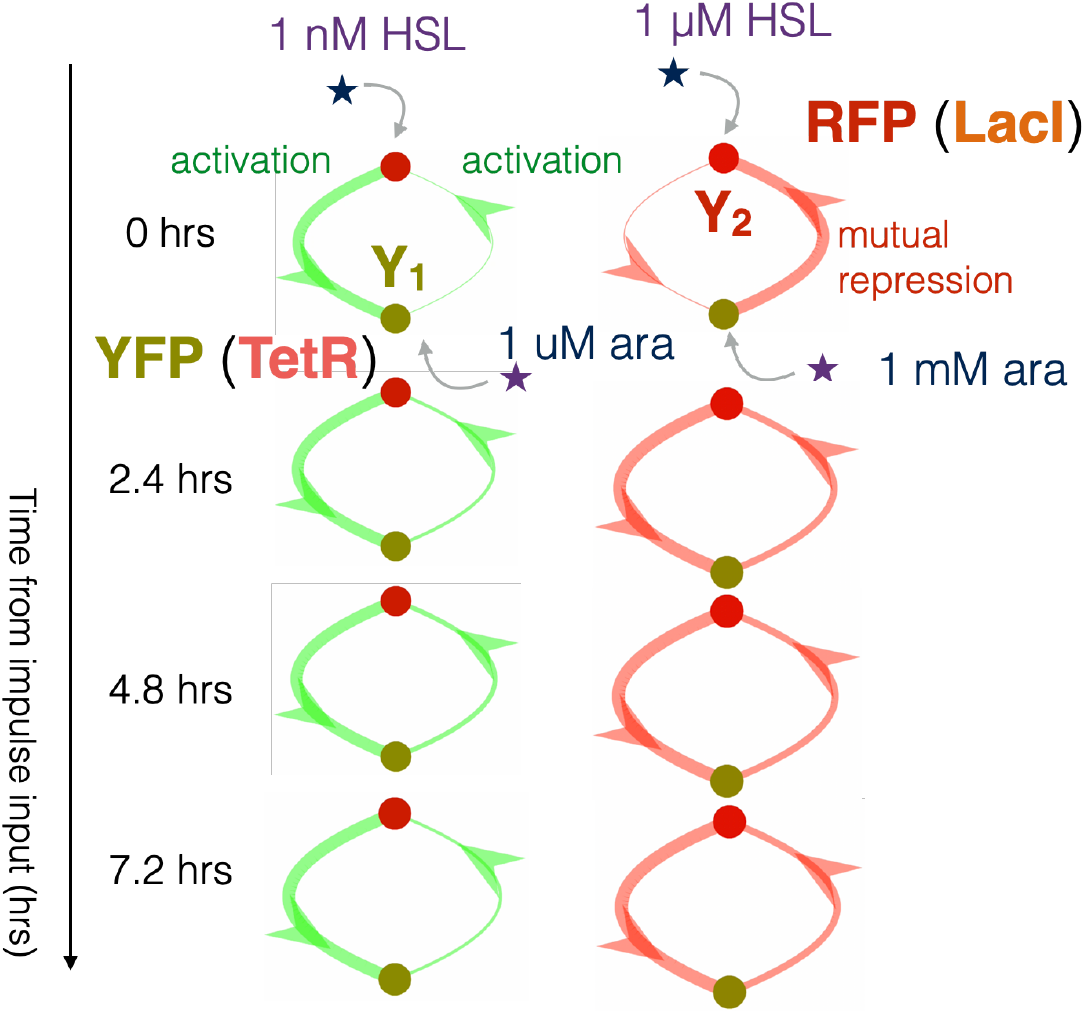
A visualization of the impulse response of the estimated convolution kernel *Q*(*t*) matrix when the event detector biocircuit is induced with low (left) versus high (right) concentrations of arabinose and HSL inducer. The width of edges in this graph coincide with the magnitude of the impulse response, while coloring is red if the sign of the impulse response for a given edge is negative (repression) and green if the given edge is positive (activation).

The reconstructed network of our transcriptional event detector reveals the functional relationship between states in the circuit at different concentration regimes. At lower concentrations of arabinose and HSL, the reconstructed transcriptional event detector network reveals functional cause of failed circuit latching. Both edges in the memory module did not repress their target promoters as intended, while the pLas-Tet promoter appears to enact a much higher gain of activated expression from HSL induction than does the activated expression of the pAra-Lac promoter in response to arabinose.

In the high gain setting, where arabinose is induced at 1 mM and HSL is induced at 1 *µ*M, we see that the memory module exhibits the proper mutually repressing motif characteristic of the genetic toggle switch up after the arrival of the HSL inducer. The repression in both edges steps up their gain as *t* approaches 4 hrs, which is roughly the time when we see a plateauing of production in the RFP signal in Figure 8C. From our reconstruction model, we can see that the edges are well balanced at the higher concentration of inducers. At the low gain of inducers, the network is completely inactive, even though the genetic sequence of the circuit is the same. This example shows that our network verification algorithm can be used to determine the conditions, or the performance envelope, under which the circuit is functioning properly. Even though the underlying model of our system is a linear approximation to a nonlinear system, we can test the system at multiple initial conditions, operating points, or equilibria, to quantify network behavior of the system locally. Taken in aggregate, these can provide a parameterized view of how the network behaves over a range of experimental conditions.

## VI. Conclusion

The dynamical structure function models the dependencies among measured states. It is a flexible representation of network structure that naturally adapts to the constraints imposed by experimental measurement. Since identifiability conditions of the dynamical structure function have been well characterized, appropriate experimental design can ensure that the process of network reconstruction produces a sensible answer.

In this work, we introduced a network reconstruction algorithm and a code base for reconstructing the dynamical structure function from data, to enable discovery and visualization of graphical relationships in a genetic circuit diagram as *time-dependent functions* rather than static, unknown weights. We proved a theorem, showing that dynamical structure functions can provide a data-driven estimate of the size of crosstalk fluctuations from an idealized model. We then illustrated these findings with numerical examples. Next, we used an *in vitro* genetic circuit, deliberately tuned with gain imbalance, to validate our algorithm on experimental data. Finally, we built a new *E. coli* based transcriptional event detector and showed how estimation of the dynamical structure reveals active and inactive network states, depending on inducer concentration. These results show how the dynamical structure function characterizes the operational or active network. They also provide a route for future study of relationships between environmental parameters, active network dynamics, and biocircuit performance.

## VII. Experimental Methods

All plasmids were constructed using either Golden Gate assembly [79] or Gibson isothermal assembly [80] in *E. coli*. Plasmids were sequence verified in JM109 cloning strains and transformed into the strain MG1655ΔLacI, provided as a courtesy by R. J. Krom and J. J. Collins. The event detector was transformed as a two-plasmid system with kanamycin and chloramphenicol selection. All *in vivo* experiments were carried out with *n* = 2 replicates using MatriPlates (Brook Life Science Systems MGB096-1-2-LG-L) 96 square-well glass bottom plates at 29*°*C in a H1 Synergy Biotek plate reader using 505/535 nm and 580/610 nm excitation/emission wavelengths. Cell density was quantified with optical density at 600 nm.

For *in vitro* experiments, all genelet repressilator reconstruction experiments were carried out at 37*°*C in a Horiba Spectrofluoremeter with 1 minute readout times, using Rhodamine Green, TYE 563 and Texas Red flourophores with 10 nm monochromator excitation and emission bands centered at 502/527, 549/563, and 585/615 nm respectively. All event detector network reconstruction reactions were performed using 500 *µ*L reaction volumes in transformed *E. coli*, grown in square well glass-bottom plates using MatriPlates (Brook Life Science Systems MGB095-1-2-LG-L) with Luria-Bertain rich media broth at 29*°*C.

## VIII. Author Contributions

E. Y. wrote the paper. E.Y, J.G., Y.Y., J. K., and R. M. M. edited drafts of the paper. E.Y. and J. K. designed and carried out experiments and processed experimental data. E.Y. performed analysis and modeling. J. G. and R. M. M. secured research funding. R. M. M. supervised the research process.

## IX. Acknowledgments

We would like to acknowledge Sean Warnick, Vipul Singhal, Shara Balakrishnan, and Anandh Swaminanthan for insightful conversations on network reconstruction algorithms. We would like to thank and acknowledge Victoria Hsiao, Ophelia Venturelli, Clarmyra Hayes, Emmanuel de los Santos, Joe Meyerowitz, and Zachary Sun for guidance with experimental techniques. This work was supported by the Engineering and Physical Sciences Research Council, the Luxembourg National Research Foundation, Air Force Office of Scientific Research, Grant FA9550-14-1-0060, the Defense Advanced Research Projects Agency, Grants HR0011-12-C-0065 and FA8750-19-2-0502, the Army Research Office Young Investigator Program, Grant W911NF-20-1-0165, the National Science Foundation, Grant 1317291, and the John and Ursula Kanel Charitable Foundation.

## X. Supplementary Information

### Experimental Methods for Circuit Preparation, Assembly, and Testing

#### A. The Repressilator Genelet Circuit

The DNA sequences for the T31, T12, T23 switch were obtained as a gift from the Winfree lab, mirroring the design identically of the repressilator genelet circuit used in [72]. Oligonucleotides were ordered with functionalized fluorophores or quenchers, corresponding to the original design of the genetic repressilator. DNA sequences were suspended in Tris-EDTA buffer for primary stock storage, while all genelet switches T12, T31, T23 added at concentrations of 75 nM, 75 nM, and 60 nM, respectively to match previous tuning experiments to balance the repressilator, with 7.5 mM working concentration of mono-NTP solution, 24 mM MgCl2, and 1x T7 expression system buffer.

DNA analogues of RNA inhibitors were added to sequester DNA activator signal from the switches as an effective step input perturbation to each node. The switches produced a RNA signal that was designed to interfere with formation of a complete promoter region of the next downstream switch in the repressilator circuit. Adding DNA served as a step perturbation to the corresponding switch. Each DNA moiety added thus had the effect of an activator. Activator DNA molecules A1, A2, and A3, each containing Iowa Black quencher were added at 75 nM, 80 nM, and 75 nM working concentration at 20 minutes from the onset of the reaction, to determine the maximum range of quenching. At 58 minutes, we added 0.7 *µ*L of pyrophosphatase, 3 *µ*L of T7 RNA Polymerase and 2.2 *µ*L of RNase H to achieve identical working concentrations as those described in [72].

#### B. The Transcriptional Event Detector Circuit

The transcriptional event detector circuit, as illustrated in Figure 7 in the main text, is composed of four distinct gene expression cassettes that define the regulatory logic of the circuit and four distinct gene expression cassettes that generate the fluorescent reporter elements of the circuit. Each gene cassette defines a transcriptional unit, with a promoter element, an RBS, a coding sequence, and a terminator sequence. Each gene cassette was cloned using a 5 part Golden Gate assembly, with a type II BsaI restriction enzyme and overhang sequences from [81], [82]. Each assembled gene cassette was cloned in JM109 *E. coli* cloning strains and sequence verified at Eurofins Genomic, by Sanger sequencing. Assembled plasmids were engineered to enable a second stage Golden Gate assembly, using the BbsI Type II restriction enzyme, and assembled to either 1) form a master regulatory logic plasmid (pEY15K), comprised of four distinct gene expression cassettes driving transcription factor or allosteric response or 2) form a master reporter plasmid comprised of four distinct reporter elements (pEY14C). Both Stage 2 assembled regulatory logic and reporter plasmids were sequence verified using Sanger sequencing (Eurofin Genomics) and transformed into MG1655ΔLacI (a gift from the Collins laboratory). The sequences for all individual plasmids and the circuit plasmids are listed in Table I.

**TABLE I:**
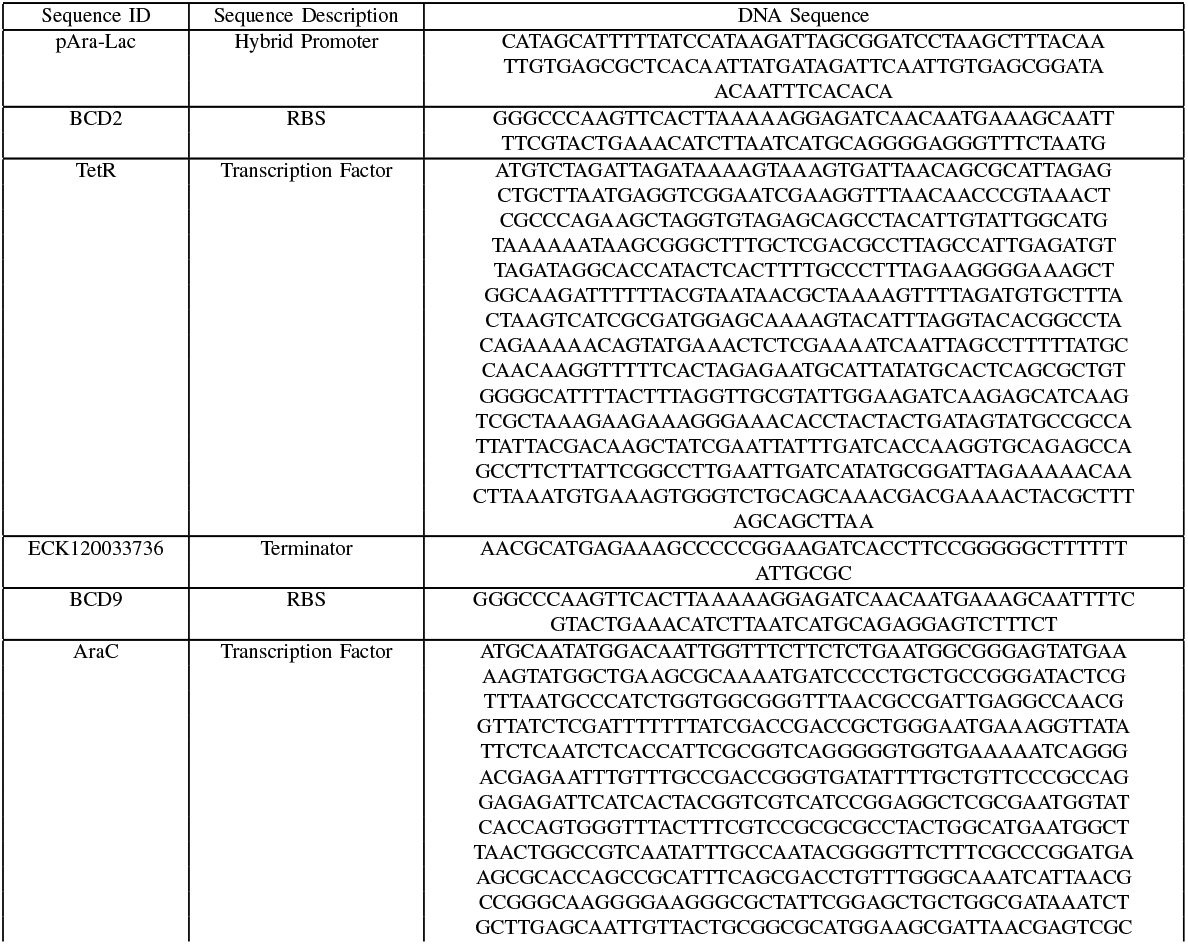
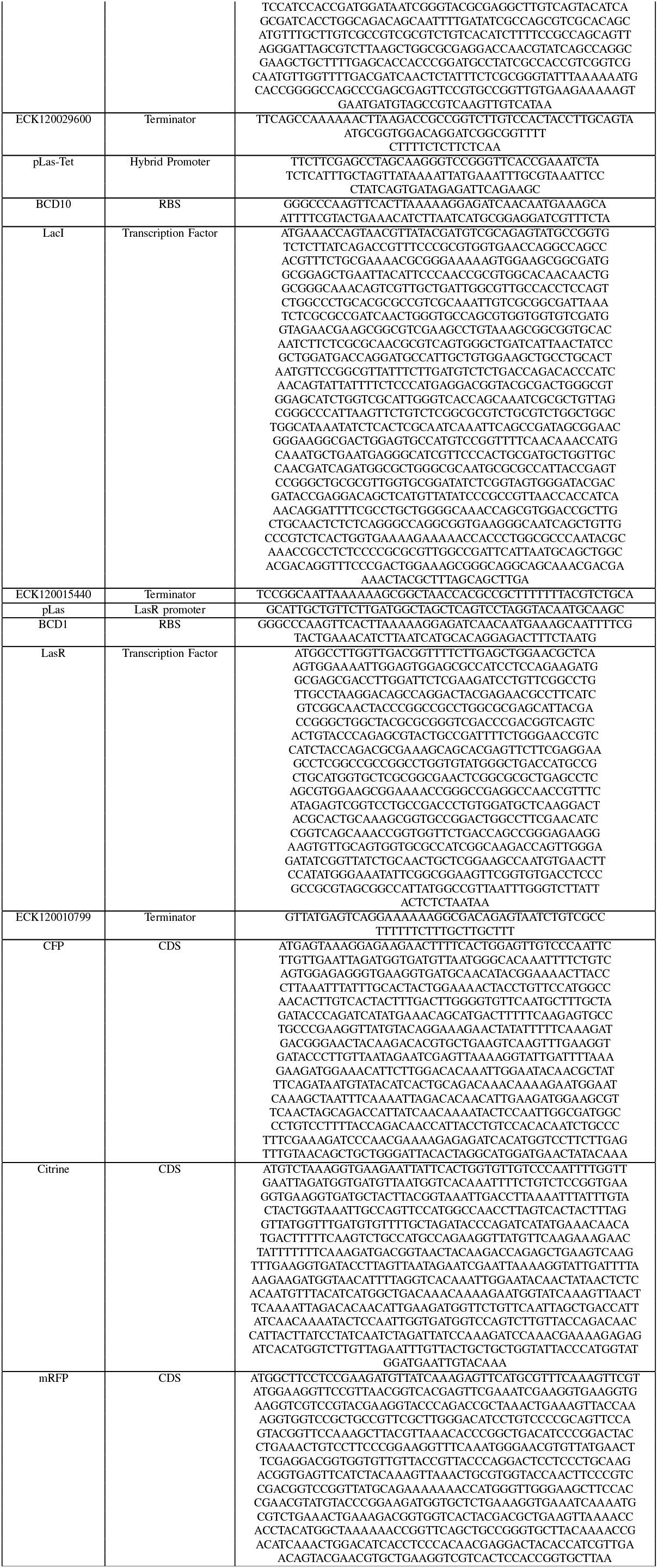
Table of genetic sequences for all parts used to make the transcriptional event detector circuit

### Sequences of Genetic Circuit Components

The sequences for all genetic components and circuits for the event detector circuit are listed in Table I. All genelet repressilator sequences are identical to the sequences used and listed in [72]. All ribosome binding site (RBS) sequences were derived from the bicistronic design (BCD) ribosome binding site library [83], while all terminator sequences were drawn from the synthetic terminator library characterized in [84].

## XI. Data Accessibility

All data files and network reconstruction code can be obtained from the GitHub repository https://github.com/YeungRepo/NetworkRecon.

## XII. Quantifying Crosstalk in Biochemical Reaction Networks

A common way that crosstalk arises in biochemical reaction networks is when species compete for commonly shared enzymes. When this occurs, the sequestration of an enzyme by one competing species makes the enzyme less accessible to other competing species. For example, when two mRNA are competing for a single ribosome, the binding of one mRNA to the ribosome during translation makes it less accessible to other mRNA. At the core of any such crosstalk is a sudden increase in the dependency of one biochemical state on another. Though enzyme loading may be a common source of crosstalk, such interactions can be modeled at a higher level of abstraction, namely how the dynamics of a given state are affected by the concentration fluctuations of other states.

Nearly every synthetic gene network implements causal dependencies among states. Often, these “designed” interactions take the form of transcription factor binding, sense-anti-sense mRNA regulation, and sequestration events. In practice, every physical system exhibits trajectories that are a mixture of the consequences of both interaction types: designed and crosstalk interactions. Throughout the course of this paper, we will denote the physical system of interest in our models as

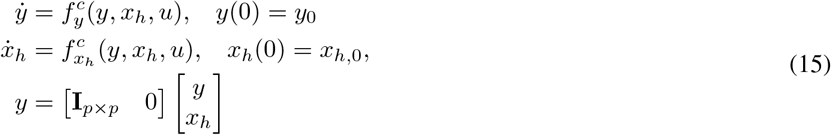

To quantify crosstalk in such systems, we can compare the dynamics of system (15) against the dynamics of a reference or alternative system that is free of crosstalk. Such a reference system will still retain the *desired* interaction dynamics and reflects the idealized model often used to design a synthetic gene network, e.g. the feed-forward loop model in Example II-A1. Moreover, it can represent the desired behavior of the system in a regime where the magnitude of crosstalk effects are supposed to be minimal or engineered in such a way that they are suppressed [7]. We write the reference system as

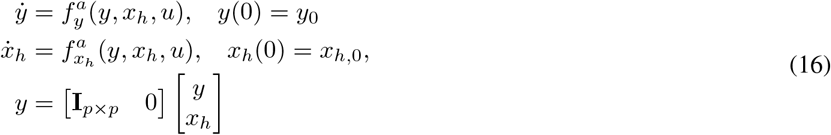

### Remark 1

For the comparison between the alternative and crosstalk system to be fair, it is important that (16) satisfies *internal equivalence* [85]. Specifically, we will suppose that any parameters or dynamics unassociated with crosstalk, e.g. interaction dynamics, catalytic reactions, or anabolic reactions with no loading effects, are held fixed. Thus, as we compare the behavior of both systems, any differences in the hidden state *x*_*h*_ or output *y* dynamics are purely due to effects of crosstalk. With the definition of an alternative system in place, it becomes possible to reason about the size of crosstalk, by comparing the dynamics of both systems. In particular, we can develop a rigorous notion for describing the amount of crosstalk arising from the difference of trajectories in both systems.

### Definition 1 (Crosstalk Trajectory)

Consider two systems, a crosstalk system and an alternative or reference system, initialized from the same initial condition *x*(0). For each initial condition *x*(0) = (*y*(0), *x*_*h*_(0)) ∈ ℝ^*n*^ and input trajectory *u*(*t*) we define the *crosstalk trajectory ζ*(*t*) as

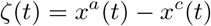

The crosstalk trajectory is a time-evolving vector that describes the deviation of the physical system (subject to crosstalk) from the reference system’s trajectory. With this notion of crosstalk, we can also make precise the concept of crosstalk between states. We note that in writing the following quantity of interest 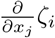, it is with a slight abuse of notation, since *ζ*_*i*_(*x*^*a*^(*t*), *x*^*c*^(*t*)). Mathematically, we are computing the *j*^th^ partial derivative of each term in 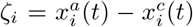. Thus, to be clear, when we write 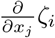, it will be implicit that we mean 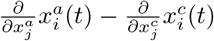.

### Definition 2 (Directed Crosstalk)

Given an initial condition of (*x*(0), *y*(0)) and input trajectory *u*(*t*) we say that a chemical species *x*_*j*_ exerts a crosstalk effect on chemical species *x*_*i*_ if the *i*^th^ component of the crosstalk trajectory *ζ*(*t*) has nonzero partial derivative

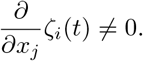

for some initial condition of (*x*(0), *y*(0)) and input trajectory *u*(*t*). In general, we will refer to 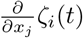 as the *crosstalk sensitivity* of *x*_*i*_ to *x*_*j*_.

Notice that the mathematical definition of crosstalk sensitivity 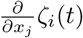 depends on the initial condition *x*_0_(*t*) and the input *u*(*t*). This dependency is consistent with the parametric sensitivity of biological function. Many genetic circuits in bacteria behave acceptably in one initial condition and for one input condition, e.g., in log-phase with an attenuated amount of a small molecule or sugar compound, but exhibit significantly different behavior when input concentrations are increased by an order of magnitude or subject to an alternate preparation method prior to the experiment. The latter imposes a state history that defines a distinct initial condition, which can drive a biological network to a highly coupled or decoupled state.

### Example 1

Consider two mRNA species *m*_1_ and *m*_2_ competing for the same degradation enzyme *D* in a physical system. For simplicity of exposition, suppose their production dynamics do not depend on each other and can be modeled as *P*_1_(*t*) and *P*_2_(*t*) respectively. The crosstalk system is given as

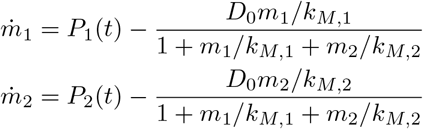

while the reference system is given as

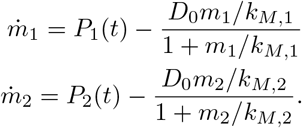

In both systems, we have supposed that time has been rescaled so that the customary parameter *k*_*cat*_ for degradation is unity. The crosstalk sensitivity of *m*_1_ and *m*_2_ (with respect to each other) are given as

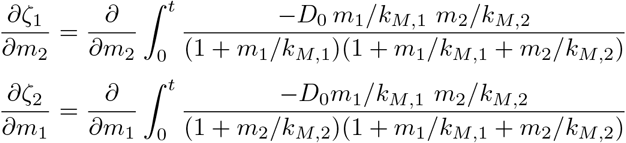

respectively. The crosstalk sensitivity between *m*_1_ and *m*_2_ is nonzero whenever *m*_1_ or *m*_2_ have non-zero initial condition.

### Remark 2

In synthetic biocircuit design, two chemical species *x*_*i*_ and *x*_*j*_ are often declared orthogonal when there is no designed interaction between them. Mathematically, in the crosstalk free system, this corresponds to

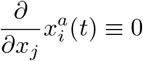

for all *x*(0) and *u*(*t*). In such a situation, *ζ*(*x*_*i*_, *x*_*j*_) ≠ 0 if and only if

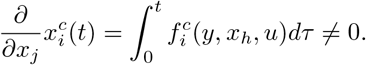

This condition is interesting in experimental settings since a computational estimate of 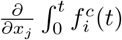 from perturbation experiments coincides with a direct estimate of the sensitivity of the crosstalk 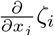. More specifically, when *x*_*i*_ and *x*_*j*_ are measured outputs of the system, we will show in the sequel that quantifying 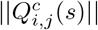 is directly related to an estimate of the crosstalk sensitivity 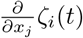 near the equilibrium point 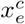.

### Remark 3

In general, estimating the crosstalk sensitivity for the nonlinear systems (15) and (16) can be challenging if either *x*_*i*_ and *x*_*j*_ are not measured directly. Firstly, if experimental data is available, it will often consist of data for the measured species *y* in the crosstalk-system, but not the reference system. Second, if only one of the species *x*_*i*_ (or none) is available for measurement, even if perturbation of *x*_*j*_ is possible, a nonlinear observer is required to estimate the trajectory of *x*_*j*_(*t*). Unless the parameters of *f*_*i*_(*x, u*) are known *a priori* (which is generally not the case), this then also requires system identification of the parameters of *f*_*c*_(*x, u*) and *f*_*a*_(*x, u*) which often results in a non-convex optimization problem.

Thus, our goal is to estimate the observed crosstalk between measured species *Y*_*i*_ and *Y*_*j*_. This crosstalk estimate will invariably include the dynamics of unmeasured chemical species (such as ATP, RNAP, untagged mRNA and protein species, DNA-protein complexes etc.). From a synthetic biology design standpoint, this is not a disadvantage, since the goal is to design a synthetic gene network with an *abstracted* circuit architecture operating reliably in the context of many unmeasured species. In any genetic circuit, there are always additional biochemical compounds that are unmeasured. Our goal is to validate that a biocircuit (e.g. an IFFL, repressilator, or a novel biocircuit) still manifests the intended network structure even in the presence of unmeasured dynamics.

### Theorem 3

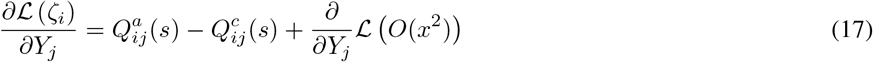

and in particular, if

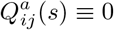

then

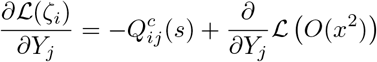

and can be estimated from input output data (*Y* (*s*), *U* (*s*)).

### Proof 1

First, notice that the Laplace transform of *ℒ* (*ζ*(*t*)) = *ℒ* (*x*^*a*^ − *x*^*c*^) ≜*X*^*a*^(*s*) − *X*^*c*^(*s*), which can be decomposed into its measured and unmeasured states

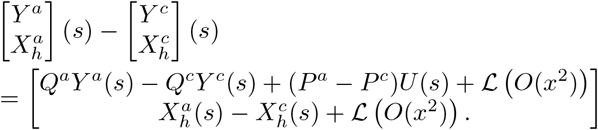

Examining the *i*^th^ component equation and taking partials along *Y*_*j*_(*s*) yields equation (17).

This result is important, since it tells us when estimating *Q*^*c*^(*s*) from experimental data will correspond to estimating crosstalk between measured states in *Y* (*s*). Since necessary and sufficient conditions for identifying *Q*(*s*) and *P* (*s*) have been already characterized [62], this provides conditions for inferring crosstalk from input-output data. For example, a sufficient condition required is that there is an input variable available to excite each measured output of the genetic network attempting to be reconstructed. This allows for the possibility that some biological states are unmeasured and unexcited, but these will be viewed as hidden states that play a role in defining the edge dynamics in *Q*^*c*^(*s*).

More generally, even if parameters for *f*^*a*^(*x, u*)(*t*) are unknown, the structure of *Q*^*a*^(*s*) can be analytically calculated (using a symbolic algebra package). For every zero entry in *Q*^*a*^(*s*) (coinciding with designed orthogonality between measured states), we can then estimate *Q*^*c*^(*s*) directly.

In practice, estimation of *Q*^*c*^(*s*) is also confounded by noise. In our analysis in this paper, we suppose that a series of filters can be applied to eliminate the noise in the data. This may not be the case for biological systems that have been characterized as inherently stochastic, e.g. single cell gene expression dynamics. In such settings, the estimated dynamical structure *Q*^*c*^(*s*) is a mixture of the process noise in the system and the crosstalk. From the standpoint of synthetic biocircuit prototyping, both are undesirable in the ultimate iteration of the biocircuit and thus need to be quantified. In this paper, we will demonstrate our theoretical and computational framework with experimental results derived from *in vitro* systems, where signal-to-noise ratios are high and the only sources of noise are measurement noise and pipetting error. For a theoretical treatment of how to reverse engineer *Q*^*c*^(*s*) in the presence of process noise or system perturbation, see [87].

An advantage of using *Q*^*c*^(*s*) to estimate the crosstalk is that we can use the *ℋ*_*∞*_ norm of 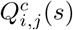 to calculate the worst-case crosstalk magnitude and *ℋ*_2_ of 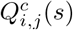 to calculate the average crosstalk across all frequencies.

## 1) Quantifying Crosstalk with Q^c^(s)

Recall the incoherent feedforward loop in subsections II-A1 and II-A2. In particular, comparing *Q*^*a*^(*s*) and *Q*^*c*^(*s*) we see that *Q*^*c*^(*s*) is a full transfer function matrix

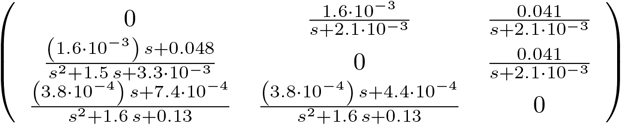

and *Q*^*a*^(*s*) is lower-triangular, reflecting the network structure of the intended IFFL. By examining the upper triangular entries in *Q*^*c*^(*s*), we can directly examine the effects of degradation crosstalk. In the lower entries of *Q*^*c*^(*s*), these crosstalk effects are confounded with the direct interactions modeled in *Q*^*a*^(*s*). Although the gain of the entries in *Q*^*c*^(*s*) are small, they nonetheless can have a significant effect on the dynamics of the IFFL.

In Figure 11 we plot the time-lapse response of *y*_2_(*t*) and *y*_3_(*t*) for varying parameter values of *k*_2,*d*_ in equation (7). The *k*_2,*d*_ parameter is a Michaelis constant that determines the effective affinity of substrate *x*_2_ in binding with *C*_0_. As *k*_2,*d*_ increases, the affinity of substrate *x*_2_ is diminished, relative to the affinity of *x*_1_ and *x*_3_. Attenuating *k*_2,*d*_ can be viewed as similar to swapping out a strong degradation marker for protease degradation with a weaker degradation marker on the species *x*_2_. In the experimental literature, there are multiple degradation markers for proteins that confer varying binding affinities to an associated protease [88]. In our simulation, we consider five potential values for *k*_2,*d*_ : 500, 1625, 2750, 3875, and 5000 *µM* corresponding to five artificial LVA markers of varying strengths for the protease ClpXP frequently used in *E. coli*.

**Fig. 11.**
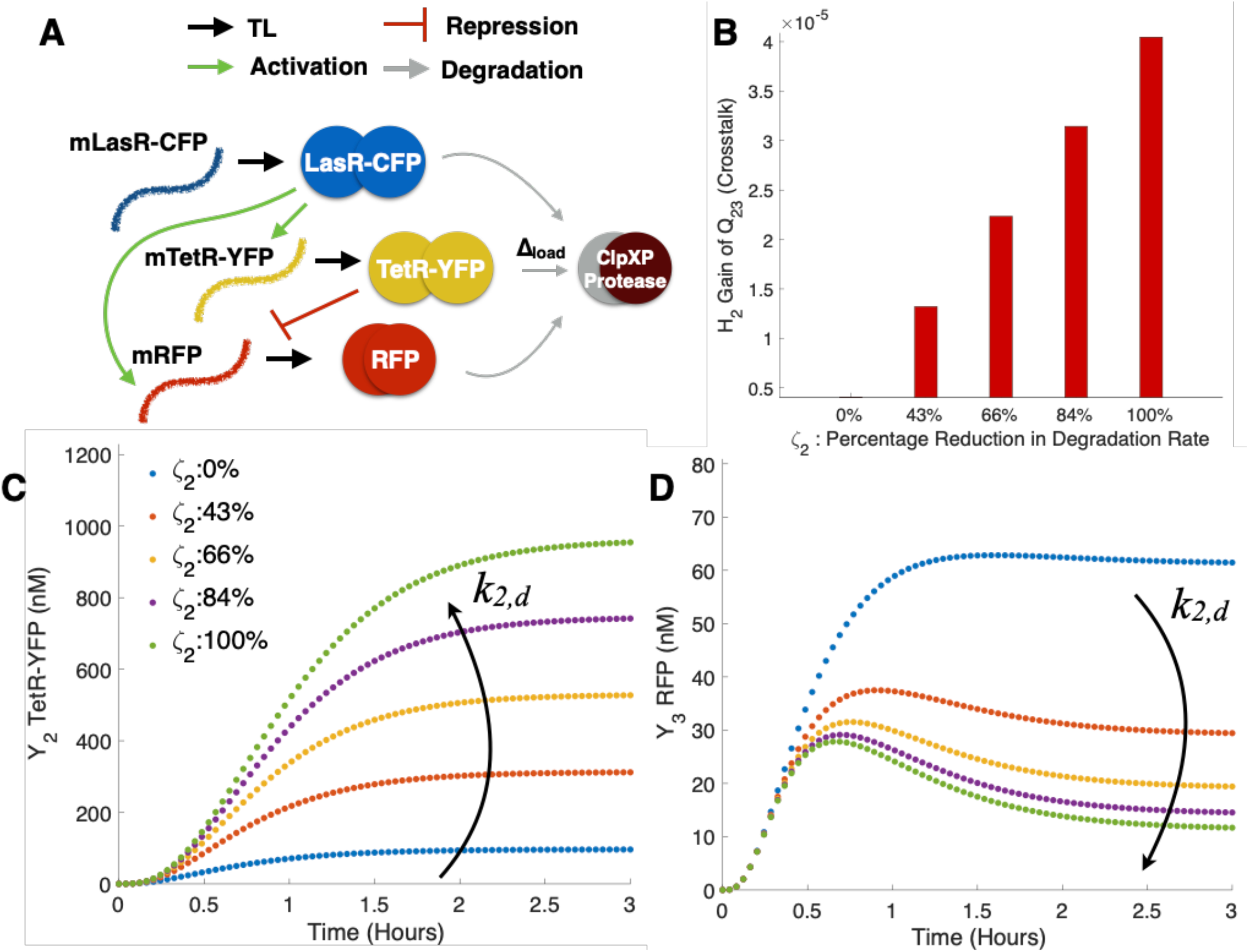
Dynamical structure functions quantify biomolecular crosstalk: **(A)** A schematic illustrating the design of this simulation example. The crosstalk and reference model of the incoherent feedforward loop from Examples II-A1 and II-A2 are simulated accordingly to satisfy internal equivalence, for varying values of *k*_2,*d*_. Standard parameters from the literature [86] were used to generate the simulation. As the size of the load Δ_*load*_ increases, the ability of the IFFL to respond with a pulse decreases.**(B)** The *ℋ*_2_ gain of 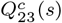 is plotted as a function of *ζ*. Notice that 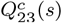 is a pure crosstalk term, since 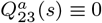. As the effective crosstalk in *ζ*_2_increases, 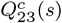 mirrors that increase, as shown in Proposition 3. **(C-D)** Time lapse responses of the incoherent feedforward loop: for each value of *k*_2,*d*_ the value of *ζ*_2_ at *t* = 3 hours is calculated and used to label curves (as percentage of maximum load). Notice the monotonic relationship between *k*_2,*d*_, *ζ* and the output responses of *Y*_2_ and *Y*_3_ (negatively monotonic).

Notice that as we decrease the affinity of *y*_2_ for ClpXP, this also coincides with an increased *ζ*_2_ crosstalk magnitude. Here, we have computed 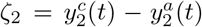. We find that |*ζ*_2_| increases as *k*_2,*d*_ increases. In Figure 11B-D, *ζ*_2_ is plotted as a percentage of maximum absolute change across all values of *k*_2,*d*_.

We see that the time-lapse response of *y*_2_(*t*) increases monotonically for all *t* as the crosstalk *ζ*_2_(*t*) increases. This is consistent with biological intuition, since an increase in competition for resource loading (an increase in *k*_2,*d*_) results in prolonged lifetimes of each individual *y*_2_ (TetR-YFP) protein. This in turn results in higher repression levels of *y*_3_ in the incoherent feedforward loop. Increased competition for ClpXP from substrates *y*_3_ and *y*_1_ have the effect of damping *y*_3_ dynamics and reinforcing the pulsatile response of the IFFL. The crosstalk in this circuit thus has the effect of effectively strengthening the negative regulation of *y*_2_ on *y*_3_, encouraging the downward transient after *t* ≊ 0.75 hours. Our network analysis shows we can improve the robustness of an IFFL’s pulse by attenuating the relative binding affinity of the repressor to its protease.

In general, crosstalk effects do not necessarily reinforce the feedback architecture of a biocircuit. This underscores the importance of having techniques for quantifying crosstalk in a synthetic gene network and validating that designed interactions are dominant over crosstalk interactions.

## Notes

### Competing Interest Statement

The authors have declared no competing interest.

### Summary of Updates

A section has been added to describe the dynamical structure estimation algorithm. The introduction and commentary throughout the paper has been revised.

